# Shared and Distinct Lipid Profiles in amygdala from Sporadic and GBA-associated Parkinson’s Diseases

**DOI:** 10.1101/2024.10.11.617800

**Authors:** Sonia S. Muñoz, Frederik R. Marlet, Jesper E. Dreier, Mesut Bilgin, Katharina Schott, Krista B. S. Neergaard, Erwan Bezard, Benjamin Dehay, Zane Jaunmuktane, Kenji Maeda, Céline Galvagnion

**Affiliations:** Department of Drug Design and Pharmacology, Faculty of Health and Medical Sciences, University of Copenhagen, 2100, Copenhagen, Denmark; Lipidomics Core Facility, Danish Cancer Institute, Copenhagen, Denmark; Univ. Bordeaux, CNRS, IMN, UMR 5293, F-33000 Bordeaux, France; Division of Neuropathology, National Hospital for Neurology and Neurosurgery, University College London Hospitals NHS Foundation Trust, London, UK; Queen Square Brain Bank for Neurological Disorders and Department of Clinical and Movement Neurosciences and Queen Square Brain Bank for Neurological Disorders, UCL, Queen Square Institute of Neurology, London, UK; Cell Death and Metabolism group, Center for Autophagy, Recycling and Disease, Danish Cancer Institute, Copenhagen, Denmark

**Keywords:** alpha-synuclein, sphingolipids, Parkinson’s Disease, amygdala, glucocerebrosidase, cholesterol

## Abstract

Parkinson’s Disease (PD) is a neurodegenerative disorder characterised by the deposition of protein-lipid inclusions, containing alpha-synuclein, neuronal cell loss and disruptions in lipid metabolism such as those associated with *GBA* mutations. *GBA* mutations are together an important genetic risk factor for PD and are associated with a decrease in glucocerebrosidase, a lysosomal glycoprotein encoded by *GBA*, increase in alpha-synuclein and changes in sphingolipids levels and composition. However, the extent of lipid metabolism disruptions associated to PD and their contributions to disease progression remain unclear.

In this study, we used a combination of biochemical and lipidomic analyses of amygdala from healthy controls (HC) and people with sporadic (sPD) or GBA-associated PD (PD-GBA) to investigate the correlation between alpha-synuclein, glucocerebrosidase and lipids.

We found extensive metabolic remodelling of brain lipids, including increased free cholesterol, diacylglycerides, sphingolipids and specific glycerophospholipids in amygdala from people with sPD and disease duration above 30 years (sPD_>30y_) and from people with PD carriers of a *GBA* risk mutation (PD-GBA_risk_) relative to HC. The levels of free cholesterol, diacylglycerides, sphingolipids and specific glycerophospholipids all correlated positively with pathological αS and negatively with GCase activity. In contrast, the levels of phosphatidylethanolamine and cardiolipin only correlated positively with GCase activity. Moreover, we observed changes in the distribution of species for sphingolipids and glycerophospholipids in opposite directions for two categories of PD cases. We found a shift from short to long sphingomyelin and ceramide and from long to short phosphatidylserine and phosphatidylethanolamine in sPD_>30y_ and PD-GBA_risk_ cases and the opposite in sPD_<10y_ and PD-GBA_severe_ cases. The relative proportion of lipid species affected in these samples all correlated with glucocerebrosidase activity and pathological alpha-synuclein levels.

Together, these findings highlight the correlation between glucocerebrosidase, pathological alpha-synuclein and lipid levels in PD. Moreover, the identified opposite changes in lipid distribution for two categories of people with sPD and PD-GBA underscore the importance of patient stratification in clinical trials aiming at reverting PD-related lipid changes.

## Introduction

Parkinson’s disease (PD) is a neurodegenerative disorder characterised by the loss of dopaminergic neurons and the deposition of protein-lipid co-assemblies called Lewy bodies (LBs) in the brain ^1,2^. These LBs are composed of protein and lipid molecules, including organelles ^2,3^. The most abundant protein is alpha-synuclein (αS), a small (14.5 kDa) intrinsically disordered protein proposed to play a role in synaptic plasticity via binding to synaptic vesicles ^4–7^. αS-membrane binding is associated with the folding of the protein into an alpha-helix^6–13^. Several studies have shown that this protein-membrane interaction is crucial not only for the proposed biological function of αS but also for modulating the formation of toxic αS amyloid fibrils ^6–13^. Indeed, αS can bind membranes of different lipid compositions but only co-assemble into mixed lipid-protein amyloid fibrils with a specific set of lipids, including negatively charged glycerophospholipids and gangliosides, at high protein-to-lipid ratios ^9,13–16^.

Disruptions in the levels of specific lipids have been observed in association with both the sporadic and familial forms of PD ^17–20^. Mutations in the *GBA* gene that encodes the lysosomal glycoprotein glucocerebrosidase (GCase) are together the most important genetic risk factor for PD ^21–23^. *GBA* mutations can be classified as severe, mild, or at risk depending on the associated severity of the phenotype ^24–27^ and were found to lead to a decrease in GCase activity in patient-derived samples, including post-mortem brain samples (i.e. cerebellum, putamen, amygdala, substantia nigra (SN) and cingulate gyrus) ^18^, cerebrospinal fluid (CSF) ^28^, fibroblasts ^29–33^ and induced pluripotent stem cell-derived neurons ^19,28,33–38^. A decrease in GCase activity due to *GBA* mutations or small molecule inhibitor (e.g. conduritol-β-epoxide) treatments was reported to be associated with increased levels of total and pathological αS and/or GCase lipid substrates, glucosylceramide (GlcCer) and glucosylsphingosine (GlcSph), in cellular models of PD and in samples from people with PD ^18^.

GCase activity has been found to decrease with ageing in the brains of healthy individuals and people with PD and to be lower in the brains of people with sporadic PD (sPD) or with PD and carriers of a *GBA* mutation (PD-GBA) relative to healthy controls (HC) ^18,39,40^. The magnitude of the decrease in GCase activity and that of the associated increase in the lipids and total/pathological αS levels were reported to vary from one brain area to another and from one cohort to another ^18^. GCase activity decreased with aging in healthy SN and putamen and in PD SN ^39,40^, and such a decrease in GCase activity was associated with increased levels of pathological αS and GlcCer only in PD SN ^40^. Studies comparing GCase activity in sPD or PD-GBA and HC gave conflicting results where GCase activity was reported to be either significantly decreased or unchanged in the cerebellum, cingulate cortex and putamen of sPD cases, SN of sPD and PD-GBA cases and frontal cortex of PD-GBA cases but to be consistently decreased in cerebellum, putamen, amygdala and cingulate cortex of PD-GBA cases or unaffected in frontal and occipital cortices in sPD ^18,19,39–43^. Decreased GCase activity was associated with unchanged GlcSph in sPD cerebellum and putamen ^39^, increased GlcCer in sPD SN ^39,40^ and increased GlcSph in sPD hippocampus and PD-GBA cerebellum, putamen, cingulate and frontal cortices ^19,39^. Finally, *GBA* mutations were associated with changes in the levels of lipids other than GCase’s substrates, including gangliosides, in specific brain areas, such as the middle temporal gyrus, cingulate gyrus and striatum ^44^ as well as disruptions in sphingolipid (SL) species distribution in anterior cingulate cortex ^45^. Despite increasing studies highlighting brain lipidome changes associated with sPD and PD-GBA, systematic analyses of the contribution of *GBA* mutations, αS, and lipids to PD and PD-GBA-related pathology are currently missing.

In this study, we used a combination of biochemical and lipidomic analyses to investigate the correlations between the levels of GCase protein and activity, pathological αS, and lipids in amygdala of HC, people with sPD and different disease duration (< 10 years (y) (sPD_<10y_), 10 – 20 y (sPD_10-20y_), 21 – 30 y (sPD_21-30y_) and > 30 y (sPD_>30y_)) and people with PD carriers of a *GBA* mutation (severe (PD-GBA_severe_), mild (PD-GBA_mild_) or risk (PD-GBA_risk_)). We found that the levels of GCase activity were decreased whereas those of pathological αS, free cholesterol (Chol), diacylglycerides (DAG), SL and specific glycerophospholipids (GPL), were increased in sPD_>30y_ and PD-GBA_risk_ amygdala compared to HC. Furthermore, the levels of Chol, DAG, SL and specific GPL all correlated positively with pathological αS and negatively with GCase activity but only for values below ca. 85% of that of HC. On the contrary, the levels of cardiolipin (CL) and phosphatidyl ethanolamine (PE) correlated positively with GCase activity over the whole value range. Moreover, our shotgun lipidomic analyses show that the hydrocarbon chain compositions of SL and GPL classes were altered in sPD and PD-GBA amygdala relative to HC but in the opposite direction for sPD_<10y_ and PD-GBA_severe_, on the one hand, and sPD_>30y_ and PD-GBA_risk_, on the other hand. Overall, we observed a shift towards shorter and more saturated phosphatidylserine (PS) and PE and towards longer and more unsaturated sphingomyelin (SM) and ceramide (Cer) in sPD_>30y_ and PD-GBA_risk_ cases and the opposite in sPD_<10y_ and PD-GBA_severe_ cases compared to HC.

## Materials and Methods

### Study design

All brain samples were obtained from Queen Square Brain Bank for Neurological Disorders (London, UK). All sPD and PD-GBA cases met the clinical and pathological criteria for advanced PD. HC controls used in the study had no clinical history of any neurological disorder and no pathological evidence of Lewy body disorder.

The sample size was determined by the available number of people with PD with *GBA* mutations, as indicated in Table 1. Amygdala tissues were dissected from fresh frozen post-mortem brains from a total of 50 individuals (15 x HC, 20 x sPD and 15 x PD-GBA (5x p.N409S, 4x p.E365K, 4x p.L483P and 2x p.T408M)).

**Table 1.**
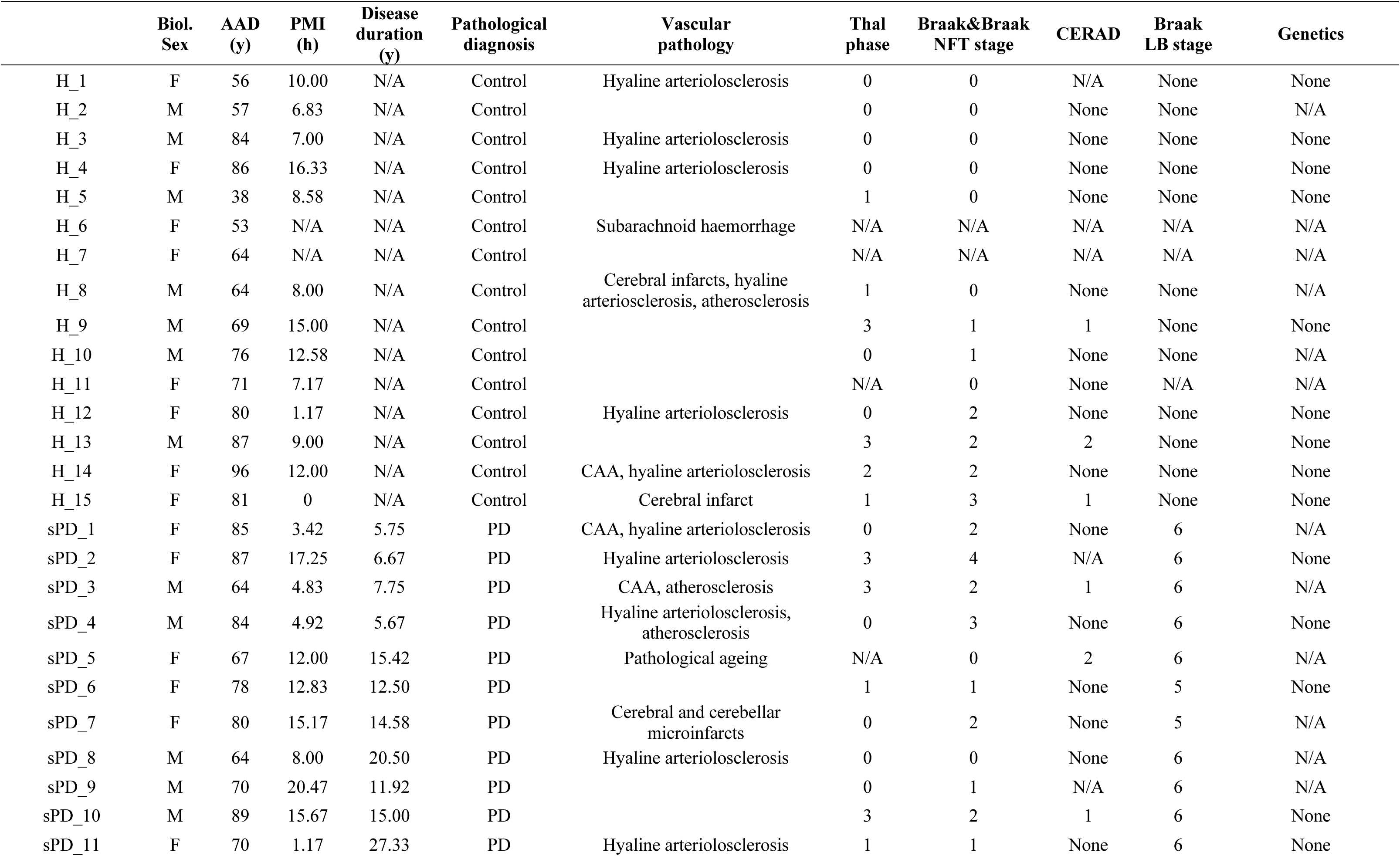
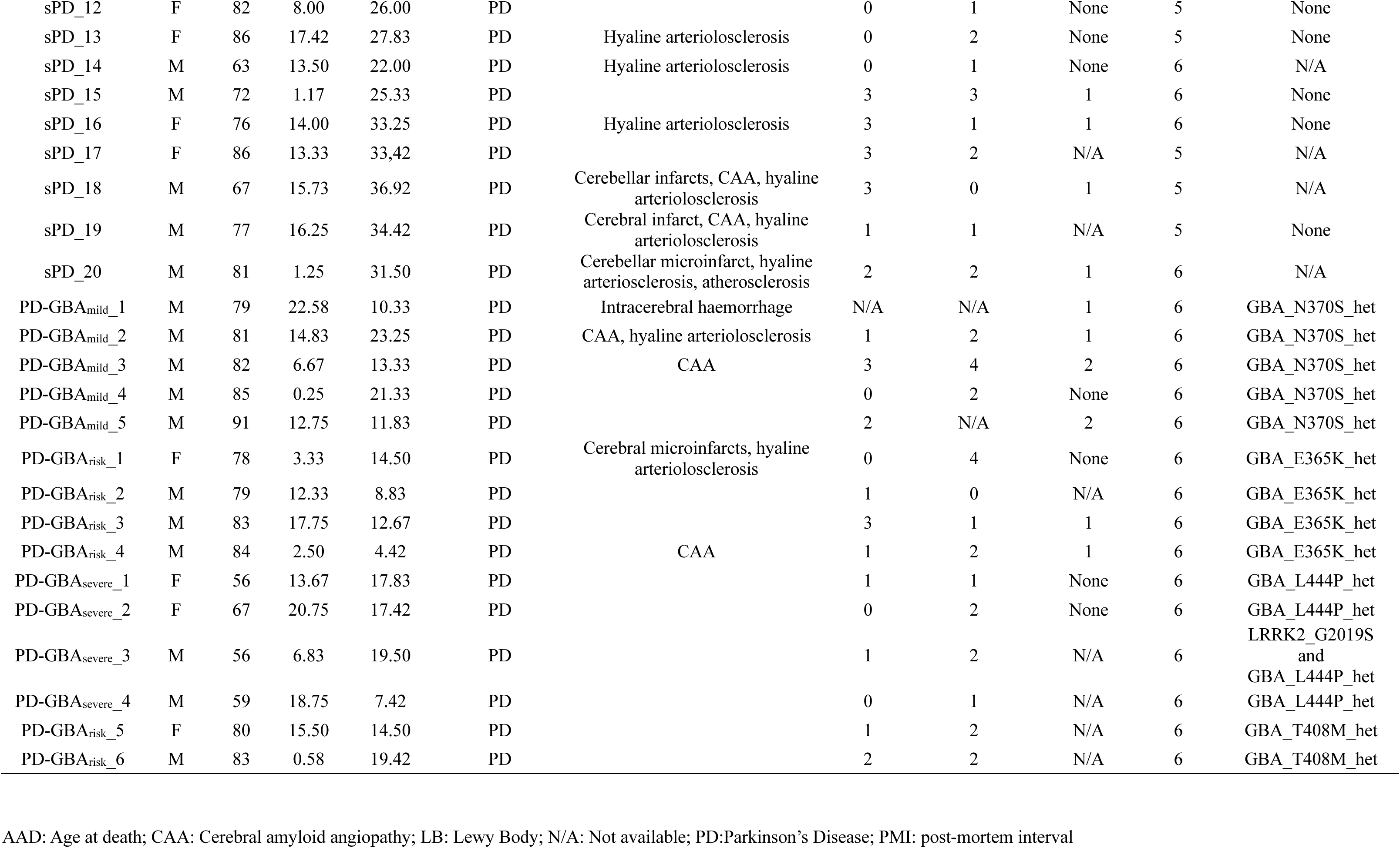
Demographic details of the patients included in the study.

### BCA Assay

Total protein content was measured in homogenate samples using the Pierce BCA Protein Assay Kit (Thermo Fisher Scientific, 23225) following the manufacturer’s protocol and every sample was run in duplicate.

### Lipidomics

Quantitative mass spectrometry-based shotgun lipidomics was performed as previously described^46^, with minor modifications. The lipid extraction was carried out at 4°C using HPLC-grade solvents in Eppendorf tubes. Homogenates (corresponding to 25 µg total proteins) were mixed with 155 mM ammonium bicarbonate (AB, Sigma-Aldrich, FL40867) to a final volume of 200 µl and sonicated for 30 min in a water-bath sonicator. Each sample was spiked with an internal lipid standard mixture (**Supplementary Table 1**). Chloroform/methanol (C/M, 2:1, v/v; 1 ml) was added, and samples were shaken for 15 min (2,000 rpm) before centrifugation (2 min, 2,000 g). The lower organic phase was transferred to a new Eppendorf tube, and two rounds of washing were performed by adding methanol (100 µl) and AB (50 µl), shaking (5 min, 2,000 rpm), and centrifuging (2 min, 2,000 g). The final organic phase was transferred to new Eppendorf tubes and evaporated for 1 h using a vacuum concentrator.

The resultant lipid film was dissolved in 100 µl C/M 1:2 (v/v) by shaking (5 min, 2,000 rpm) and centrifuged (15 min, 15,000 g) to remove insoluble particles. Extracted lipids were then mixed with positive and negative ionisation solvents (13.3 mM AB in 2-propanol or 0.2% (v/v) methylamine in C/M 1:5 (v/v), respectively) and analysed in the positive and negative ion modes on quadrupole-Orbitrap mass spectrometer Q Exactive (Thermo Fisher Scientific) equipped with TriVersa NanoMate (Advion Biosciences) for automated and direct nanoelectrospray infusion.

Lipids were identified based on predefined criteria for each lipid class (**Supplementary table 2**). Lipid identification and extraction of precursor- and fragment-ion intensities were carried out using LipidXplorer version 1.2.4^47^, and lipid quantities in samples were calculated using the R-based in-house software LipidQ (https://github.com/ELELAB/lipidQ).

To enable specific quantification of galactosylceramide (GalCer) and GlcCer species, an additional LC-MS-based analysis was performed. For this, 50 µl of lipid extracts were transferred into glass vials, evaporated for 30 min, and reconstituted in 50 µl acetonitrile.

### LC-MS Analysis of Galactosylceramide and Glycosylceramide

GalCer and GlcCer species were analysed using liquid chromatography-mass spectrometry (LC-MS). Chromatographic separation was performed under isocratic conditions at a flow rate of 15 µL/min. Mobile phase A consisted of 95% acetonitrile (ACN), 2.5% methanol (MeOH), 2.5% water (H₂O), 0.5% formic acid (FA), and 5 mM ammonium formate. Mobile phase B consisted of 100% ACN.

The LC system was coupled to a quadrupole-Orbitrap mass spectrometer (Q Exactive, Thermo Fisher Scientific) via an UltiMate 3000 HPLC system (Thermo Fisher Scientific). Samples were analysed in positive ion mode with mass spectrometric detection performed with a scan range of m/z 500–900. The instrument was operated at a resolution of 70,000 at m/z 200, with two microscans per scan event. The maximum injection time was set to 128 milliseconds, and the automatic gain control (AGC) target was 5 × 10⁵.

We considered a lipid specie to be detected, and thereby to be included in the processing of the full dataset, if its level was a nonzero value in more than half of the total number of samples per group.

### Western blot

Protein content of homogenate samples was measured using the Pierce TM BCA Protein Assay Kit (Thermo Fisher Scientific, 23225). Samples of 40 μg were electrophoresed on a NuPage 4%-12% Bis-Tris Protein gel (Thermo Fisher Scientific) in a randomised order. Proteins were transferred to a PVDF membrane (0.2 μm, Thermo Fisher Scientific, 88520), blocked in 5% w/v milk (2 h) and treated with primary and secondary antibodies. Antibody binding was detected using an ECL chemiluminescence kit (Bio-Rad, 1705061). In order to detect multiple proteins for the same set of samples, the membrane was stained multiple times. Firstly, the full membrane was stained for pSer-αS and imaged, then the membrane was reactivated in methanol, stripped with acetic buffer (0.015 % w/v glycine, 0.001 % w/v SDS and 1% v/v Tween-20, pH 2.2), blocked in 5% w/v milk (2 h), reprobed with next antibody and imaged. Second reprobe was for total αS and third for GCase (only top part of membrane stained, >50 kDa) and β-act (only bottom part of membrane stained, <50 kDa). Western blot was performed once per sample.

The following antibodies were used: pSer129-αS (CST, 23706, 1:1000), α-synuclein (BD Biosciences, 610787, 1:500), β-actin (ABclonal, AC026, 1:100.000), GBA (Abcam, ab55080, 1:1000), anti-mouse (Abcam, ab205719, 1:5000) and anti-rabbit (Abcam, ab6721, 1:5000).

### GCase activity assay

Samples of brain homogenates were diluted to a total protein concentration of 0.1 mg/ml for PD samples and 0.06 mg/mL for HC samples in reaction buffer (0.1 M sodium citrate, 0.1% Triton X-100, 6.5 M sodium taurocholate, 2.5 mM 4-methylumbelliferyl β-D-glucopyranoside) and incubated in non-binding plates (Corning 3881) under shaking conditions (300 rpm) at 37°C.

Solutions with varying concentrations of 4-methylumbelliferone, from 0 to 50 µM, also diluted in reaction buffer, were analysed under same conditions to convert fluorescence units into µM of 4-methylumbelliferone. Fluorescence was measured using CLARIOstar plate reader (BMG Labtech, Aylesbury, UK) with excitation/emission filters of 360-20/450-10 nm. The GCase activity was calculated using following equation:

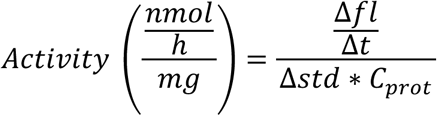

Where Δ*fl*/Δ*t* is the change in fluorescence per hour, Δ*std* is the change in fluorescence (fluorescence unit) per concentration 4-methylumbelliferone (µM) and C_prot_ is the total protein concentration of the solution, 0.05 mg/ml. Every sample was run in quadruplicate.

### Statistics

Statistical analysis and graph visualisation were performed using Prism 9 software (GraphPad) and Python. Statistical significance, when comparing the mean of two groups, was determined using unpaired t-test. Comparisons among three groups or more was performed using One-way ANOVA and multiple *t*-test comparisons (Šídák’s multiple comparisons test). Comparisons with a p-value below 0.05 were considered statistically significant. Data were shown as mean ± SEM for all results. Abundance of lipid classes and species were illustrated as Pearson r coefficients in heat maps and effects achieving *P* < 0.05 were interpreted as statistically significant. Correlation graphs were analysed using simple linear regression, with *P < 0.05* interpreted as a significant correlation between the two tested variables.

To visualise clustering in our data we used the tSNE model from scikitlearn^47^ with perplexity set to 15 and other settings set to default.

To detect when there was a break in the correlation between GCase activity and lipid abundance we used the python package PieceWise Linear Function (PWLF) ^48^. For lipid classes with a break point detected, the break point value was extracted, and the average value for all break points was used to split the data into high and low GCase activity for each lipid class.

To estimate the relationship between lipid abundance and pSer-αS and GCase activity, respectively, we applied a multiple linear regression model using the statsmodel package in python^49^. The correlation model describes the abundance of each lipid class in pmol/ 25 µg protein or the abundance of each lipid specie in percentage of lipid class as a function of both pSer-αS and GCase activity:

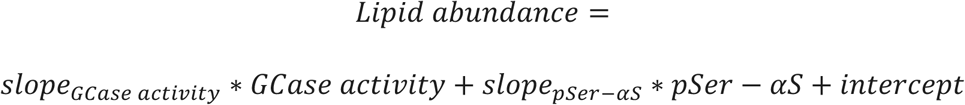

where the slopes are reported as the correlation coefficients.

The lipid data was standardised to enable comparison between the slopes of different lipid classes in the heatmaps and volcano plots, using scikitlearn standard scaler^47^.

Statistical significance of the resulting coefficients was tested using a two-tailed students t-test.

## Results

### Decreased GCase activity and increased pathological αS in sPD>30y and PD-GBArisk amygdala

To investigate lipid changes associated with PD and αS pathology in the amygdala, we examined four groups of people with sPD and different disease duration: < 10 y (sPD_<10y_, *n* = 4), 10 – 20 y (sPD_10-20y_, *n* = 6), 21 – 30 y (sPD_21-30y_, *n* = 5) and > 30 y (sPD_>30y_, *n* = 5), three groups of people with PD carriers of a *GBA* mutation (PD-GBA) categorised as severe (PD-GBA_severe_, *n* = 4, p.L483P), mild (PD-GBA_mild_, *n* = 5, p.N409S) or risk (PD-GBA_risk_, *n* = 6, p.E365K and p.T408M), as described previously ^24–27^, and one group of healthy controls (HC, *n* = 15) (see **Table 1** for demographic, clinical and genetic data). Furthermore, we selected the amygdala because of its high αS pathology burden in the majority of PD cases, contributing to PD non-motor symptoms ^50–52^.

We first determined the levels of GCase activity and protein and total and pathological αS in HC, sPD and PD-GBA homogenates and found that they correlate with neither the age at death (AAD) nor the post-mortem intervals (PMI) (**Supplementary Fig. 1**) and do not significantly differ between female and male cases for each group (HC, sPD and PD-GBA) (**Supplementary Fig. 2**). GCase activity did not correlate with disease duration in sPD homogenates (**Supplementary Fig. 2a**), but its levels were the lowest in sPD_>30y_ cases (**Fig. 1a**). We observed a trend towards lower values of GCase activity in PD-GBA_mild_ and PD-GBA_risk_ cases compared to HC (**Fig. 1a**). GCase protein correlated with GCase activity but not with disease duration (**Fig. 1c**) and was higher in sPD_10-20y_, sPD_21-30y_ and PD-GBA_risk_ than in HC (**Fig. 1b**). The level of pathological αS (pSer-αS) was semi-quantified using western-blot (WB) and defined for each sample as the ratio between the levels of total αS phosphorylated on Ser-129 (D1R1R) and that of total αS (Syn 1) (**Supplementary Fig. 3**).

**Figure 1:**
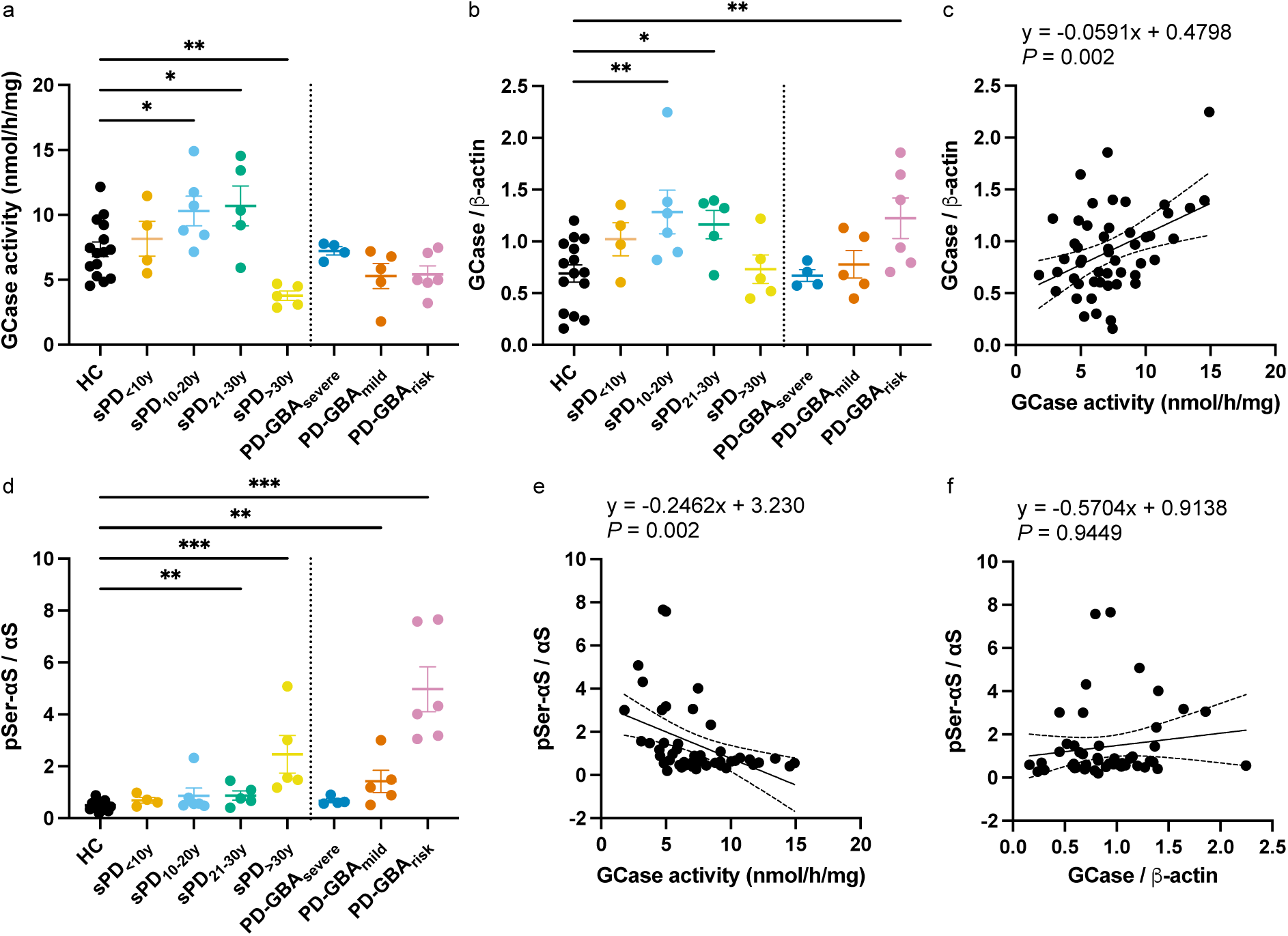
GCase protein, GCase activity and pSer-αS are altered in sPD and PD-GBA cases relative to HC. **(a,b,d)**. GCase activity (**a**) and GCase protein (**b**) and pSer-αS (**d**) measured in amygdala homogenates from HC (*n* = 15), people with sPD with disease duration below 10 y (sPD_<10y,_ *n* = 4), 10-20 y (sPD_10-20y,_ *n* = 6), 20-30 y (sPD_21-30y,_ *n* = 5) and above 30y (sPD_>30y,_ *n* = 5) and people with PD carriers of a *GBA1* mutation characterised as severe (PD-GBA_severe,_ *n* = 4), mild (PD-GBA_mild,_ *n* = 5) or risk (PD-GBA_risk,_ *n* = 6). Analysis: t-test comparison between the mean of HC vs that of each of the seven PD groups. \**P* < 0.05, \*\**P* < 0.01, \*\*\**P* < 0.001. **(c,e)** Variation of the levels of GCase protein (**c**) and pSer-αS (**e**) with those of GCase activity. (**f**) Variation of pSer-αS with GCase protein. The solid line on **c,e,f** shows simple linear regression to test the association between the levels of GCase protein (**c**) and pSer-αS (**e**) with those of GCase activity or between the levels of pSer-αS with those of GCase protein (**f**).

The levels of pSer-αS correlated with disease duration for sPD cases only (**Supplementary Fig. 4c**) and were significantly higher in sPD_21-30y_, sPD_>30y_ and PD-GBA_mild_ PD-GBA_risk_ amygdala compared to HC (**Fig. 1e,f**). Moreover, the levels of pSer-αS correlated negatively with GCase activity levels (**Fig. 1e**) but not with the levels of GCase protein (**Fig. 1f**).

### Increased levels of SL, DAG, cholesterol, and specific GPL in sPD>30y and PD-GBArisk amygdala

We then investigated the lipidome of these amygdala homogenates using mass spectrometry-based shotgun lipidomics. 330 lipid species were detected, and an average of 17,640 (± 6,538) pmol per 25 μg protein was present in healthy homogenates. Healthy homogenates were composed of the following lipid categories: 47% sterols (inc. 47% Chol + 0.03% cholesteryl ester (CE)), 45% GPL, whose main classes were phosphatidylcholine (PC) (25%), ether phosphatidylethanolamine (PE O-) (9.2%), PE (4%), PS (2.5%), ether phosphatidylcholine (PC O-) (1.1%), 7.8% SL (including SM (5.2%), Hexosylceramide (HexCer) (1.5%), Cer (0.4%), sulfatides (SHexCer) (0.4%), and diHexCer (0.2%)), and 0.2% DAG (**Fig. 2a**). Lipid levels were found not to correlate with AAD (except for LPS) and PMI and the lipidome did not vary between male and female cases for each group (HC, sPD, PD-GBA_severe_ and PD-GBA_risk_) (**Supplementary Table S3** and **Supplementary Fig. 2**). We then used t-distributed stochastic neighbour embedding (t-SNE) to visualise the amygdala lipidome variation based on the 330 lipid species detected across the eight different groups, i.e. HC, sPD_<10y_, sPD_10-20y_, sPD_21-30y_, sPD_>30y_, PD-GBA_severe_, PD-GBA_mild_, PD-GBA_risk_ (**Fig. 2c**).

**Figure 2:**
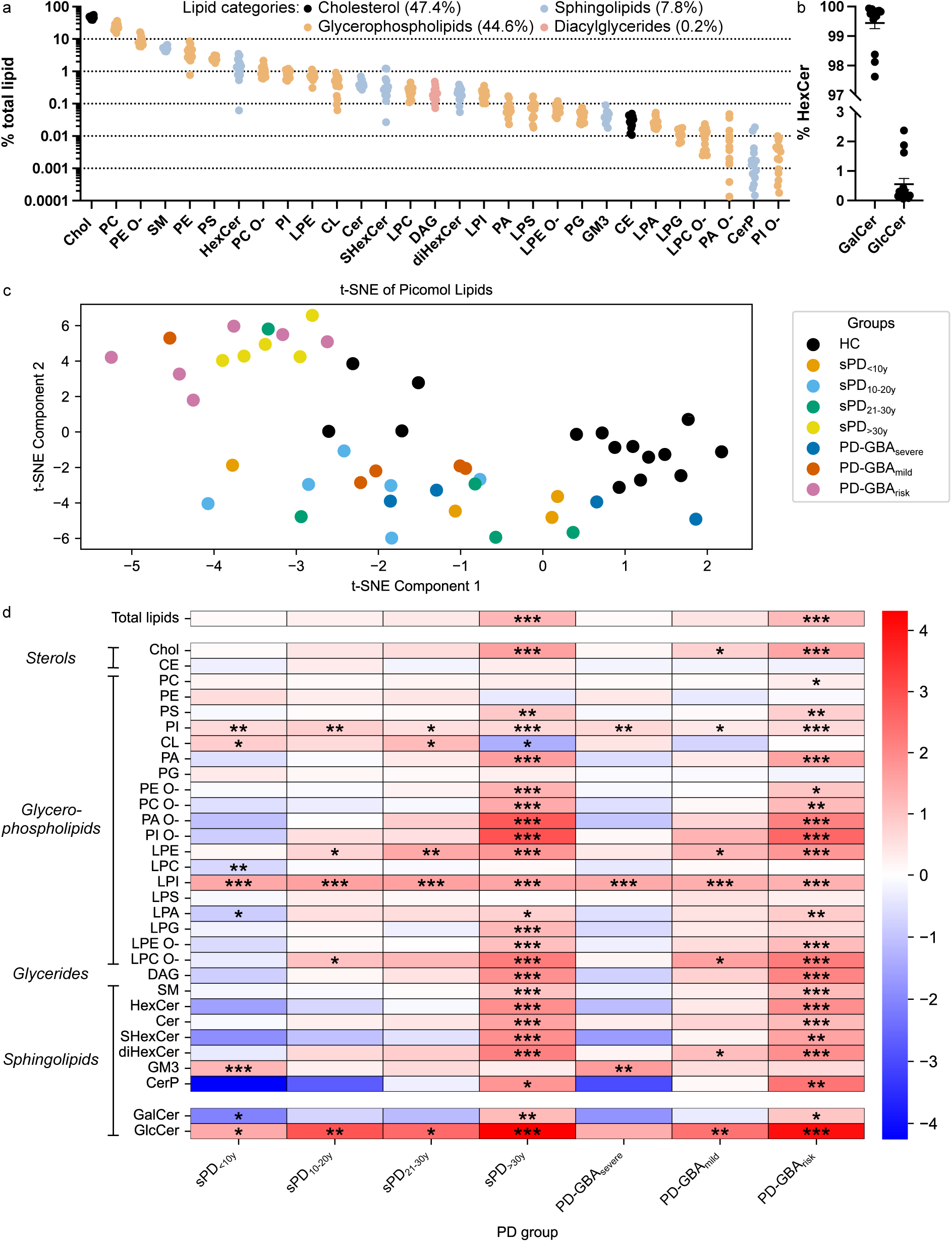
Changes in the levels of lipid classes in PD amygdala compared to HC. **(a)** Levels of each lipid classes detected in healthy amygdala expressed as percentage of total lipid. **(b)** Levels of GlcCer and GalCer in HexCer expressed in percentage of total HexCer. **(c)** t-SNE visualisation of the lipid data obtained for the HC (black), sPD_<10y_ (orange), sPD_10-20y_ (light blue), sPD_21-30y_ (green), sPD_>30y_ (yellow), PD-GBA_severe_ (dark blue), PD-GBA_mild_ (red) and PD-GBA_risk_ (pink). **(d)** Heat maps showing the log_2_FC in the levels of each lipid class in sPD and PD-GBA cases relative to that in HC. The stars indicate the p-value for the t-test comparison between the mean of the class level in sPD or PD-GBA cases and that in HC. \**P* < 0.05, \*\**P* < 0.01, \*\*\**P* < 0.001.

We observed that the eight groups formed three clusters: (i) HC, (ii) sPD_>30y_ and PD-GBA_risk_ and (iii) sPD_<10y_, sPD_10-20y_, sPD_21-30y_, sPD_>30y_, PD-GBA_severe_, PD-GBA_mild_, suggesting similarities in the lipidome of the amygdala within each cluster. Indeed, the heat map showing the log 2-fold change in the level of each lipid class (e.g. PC, PE, HexCer) in each of the seven PD groups relative to those in HC shows that the same lipid classes were increased in sPD_>30y_ and PD-GBA_risk_ cases (**Fig. 2d**). As expected for samples with decreased GCase activity levels, we observed increased levels of HexCer (composed of GlcCer, GCase lipid substrate, and GalCer) in sPD_>30y_ and PD-GBA_risk_ cases relative to those in HC (**Fig. 2d**). We performed Liquid Chromatography Mass Spectrometry (LC-MS) analyses of HexCer to separate and quantify GlcCer and GalCer and found that HexCer was composed of ca. 99% GalCer and ca. 1% GlcCer (**Fig. 2b**), in agreement with previously reported GlcCer/GalCer distribution of brain HexCer ^53^. As observed for HexCer, the levels of both GlcCer and GalCer were increased in sPD_>30y_ and PD-GBA_risk_ cases (**Fig. 2c**). The increase in lipid levels observed in sPD_>30y_ and PD-GBA_risk_ amygdala was not limited to HexCer (GlcCer and GalCer) but extended to all other SL classes, i.e. SM, Cer, diHexCer, ShexCer, GM3 and CerP. On the other hand, while Chol followed the same trend as that observed for SLs, GPL classes were either unchanged or increased, with the major GPL classes PC and PE being unchanged while PS, PI, PC O- and PE O- were increased in sPD_>30y_ and PD-GBA_risk_ cases relative to those in HC. Finally, we found that the total lipid levels were also increased in sPD_>30y_ and PD-GBA_risk_ cases relative to HC (**Fig. 2c**).

### Chol, DAG, SL and specific GPL correlate significantly with GCase activity and pathological αS

We then tested the correlation between the levels of lipid classes and disease duration as well as the levels of GCase activity, GCase protein, and pSer-αS measured in healthy and PD amygdala homogenates. We found that only PI levels correlated with GCase protein (**Supplementary Table S3**). However, the levels of lipid classes that were increased in sPD_>30y_ and PD-GBA_risk_ cases compared to HC all correlated with GCase activity and pSer-αS (**Fig. 3**). In particular, the levels of Chol, DAG, all SL (except GM3), PS, PA, all ether GPL correlated negatively with GCase activity only for GCase activity values below ca. 6.2 nmol/h/mg protein (ca. 85% of GCase activity in healthy amygdala), as determined from the fits of the data to a piecewise linear function (see Materials and methods for more details) (**Supplementary Fig. 4**), and positively with pSer-αS for the entire range of values (**Fig. 3c-e**). On the contrary, the levels of CL and PE correlated positively with GCase activity for the entire value range (**Fig. 3a,b,e**).

**Figure 3:**
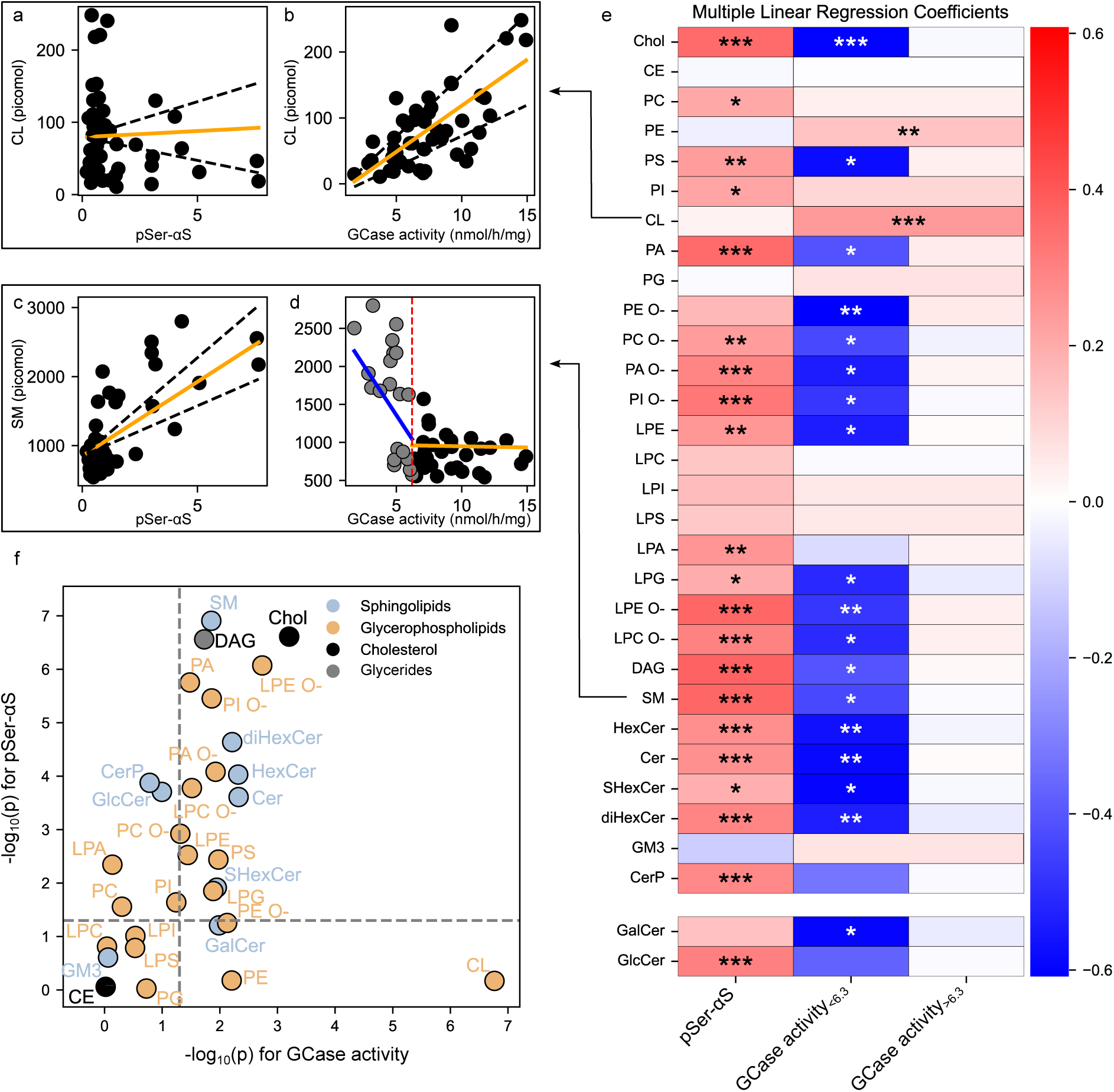
Lipid classes correlate with GCase activity and/or pSer-αS. **(a-d)** Variation in the levels of CL (**a,b**) or SM (**c,d**) with pSer-αS (**a,c**) or GCase activity (**b,d**). (**e**) Heat map showing the multiple linear regression coefficients for each lipid classes and pSer-αS, low and high GCase activity (See **Supplementary Fig. 5 and 6** for the multiple linear regression of the lipid class levels). The lipid levels used in the model were standardised for visualisation using the standard scaler function from the scikitlearn library. (**f**) Log-log plot showing the correlation between lipid class with GCase activity and/or pSer-αS. P-values: t-test of the correlation coefficients from the multiple linear regression model. \**P* < 0.05, \*\**P* < 0.01, \*\*\**P* < 0.001.

Finally, we used multiple linear regression to test the correlation between the levels of each lipid class and GCase activity or pSer-αS by adjusting for the contribution of pSer-αS or GCase, respectively (**Fig. 3f**). We found that the levels of CL and PE mainly correlated with GCase activity, whereas those of all SL (except GM3), Chol, DAG, PS, PA and ether GPL correlated with both GCase activity and pSer-αS.

All together, these results suggest that the levels of lipid classes altered in sPD_>30y_ and PD-GBA_risk_, i.e. Chol, DAG, SL and specific GPL, all correlate with either GCase activity or pSer-αS or with both. The levels of CL and PE (two mitochondrially synthetised lipids) correlate differently with GCase activity and pSer-αS than Chol, DAG, SL and GPL suggesting that the lipidome of different organelles might be affected differently by changes in GCase activity and pSer-αS levels.

### Opposite changes in GPL and SL species distribution in sPD>30y/PD-GBArisk and sPD<10y/PD-GBAsevere

Each lipid class is composed of species with varying number of total number of carbons and double bonds in their hydrocarbon chains; the sphingoid bases and N-acyl group for SL and acyl and alkyl chains for GPL. **Fig. 4,5** and **Supplementary Fig. 8 – 12** show the species distribution for the most abundant SL and GPL, where each lipid species is indicated in the format [number of carbons]:[number of double bonds];[number of OH groups] or [number of carbons]:[number of double bonds] in the hydrocarbon chains for SL or GPL, respectively. Species are expressed as relative proportion, i.e. mol%, within each SL or GPL class. We compared the species distribution of SL and GPL for each sPD disease duration group and PD-GBA groups to that of HC. We found that the same lipid species were affected in the different PD groups when compared to HC but in opposite directions for the following two categories of people with PD: (i) sPD_>30y_ and PD-GBA_risk_ on the one hand and (ii) sPD_<10y_ and PD-GBA_severe_ on the other hand (**Fig. 4, 5 and Supplementary Fig. 8 – 12**).

**Figure 4:**
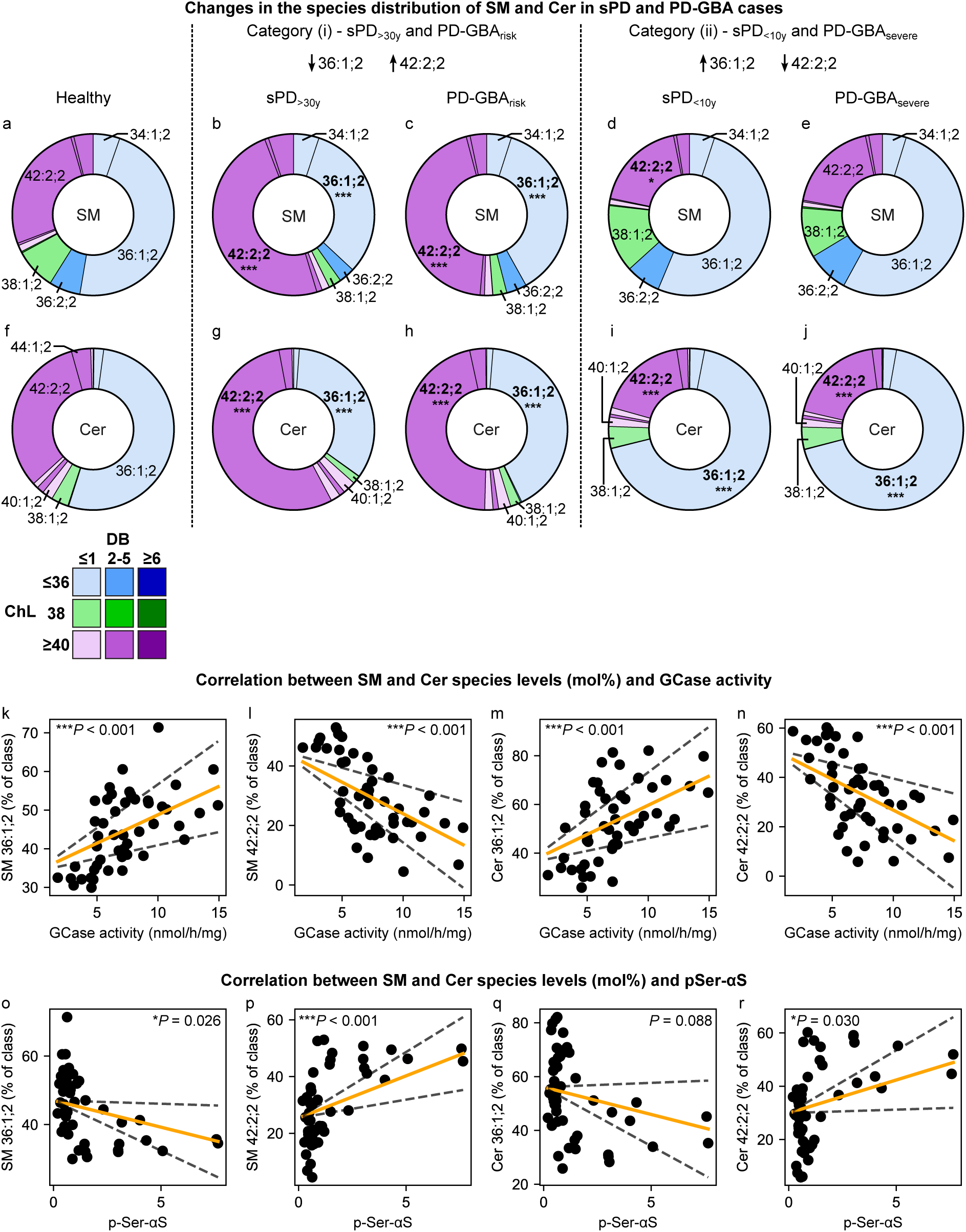
The levels of specific SL species are affected in sPD and PD-GBA amygdala homogenates and correlate with GCase activity and p-Ser-αS. (**a-j**) SM (**a-e**) and Cer (**f-j**) species distribution in amygdala from HC (**a,f**), people with sPD_>30y_ (**b,g**) and PD-GBA_risk_ (**c,h**) (category (i)), sPD_<10y_ (**d,i**) and PD-GBA_severe_ (**e,j**) (category (ii)). Analysis: 2-way anova with individual comparison mean HC vs each mean of PD group for each species. **(k-r)** Changes in the levels of SM 36:1;2 (**k,o**), SM 42:2;2 (**l,p**), Cer 36:1;2 (**m,q**) and Cer 42:2;2 (**n,r**) expressed as mol% of lipid class with GCase activity (**k-n**) or pSer-αS (**o-r**). Analysis: Multiple linear regression of the lipid data, GCase activity and pSer-αS. The black stippled lines represent the 95% confidence interval of the regression model to the data. P-values: t-test of the correlation coefficient of the fitted parameter (GCase activity (**k-n**) or pSer-αS (**o-r**)). \**P* < 0.05, \*\**P* < 0.01, \*\*\**P* < 0.001.

**Figure 5:**
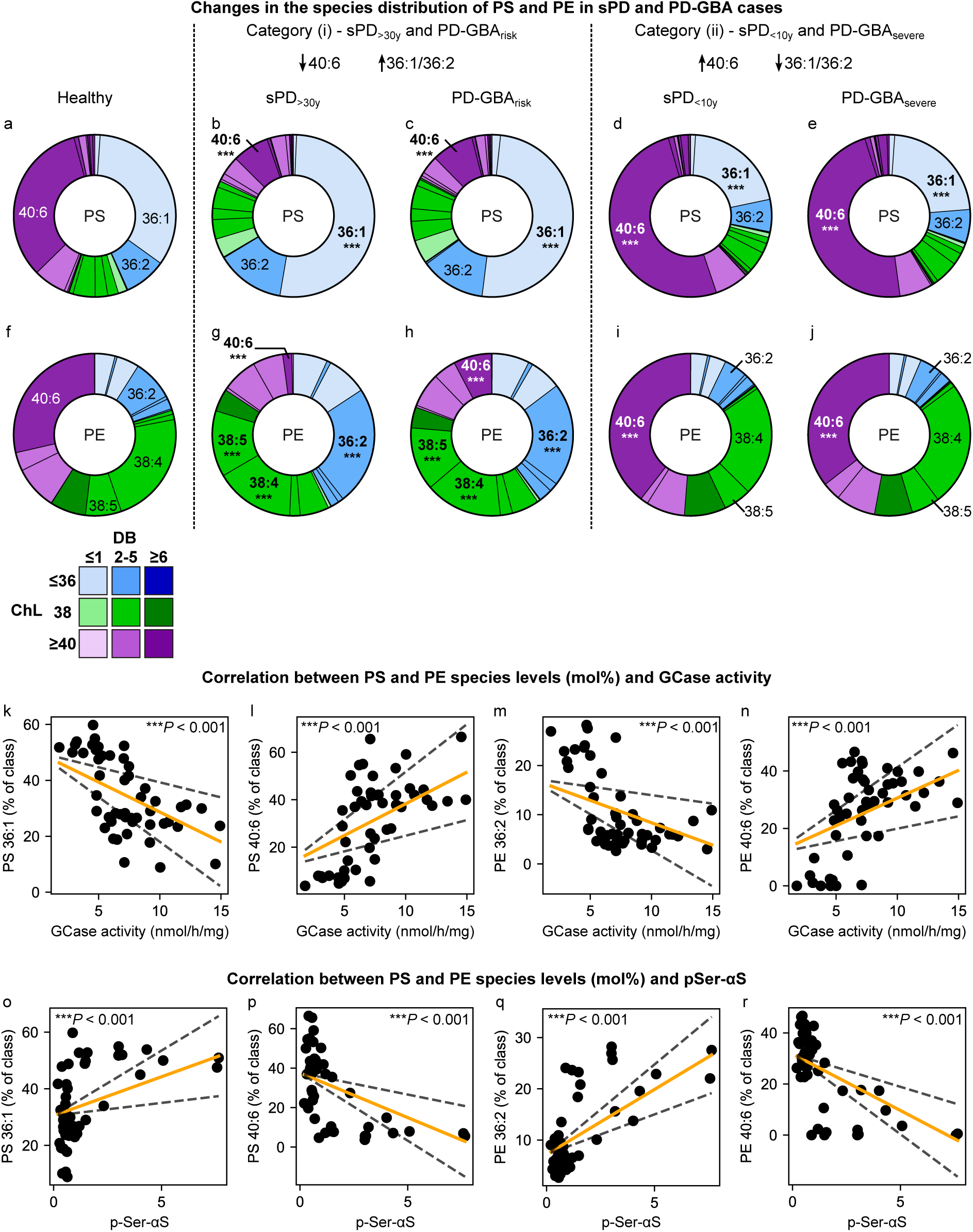
The relative levels of specific GPL species are affected in sPD and PD-GBA amygdala homogenates and correlate with GCase activity and p-Ser-αS. (**a-j**) PS **(a**-**e)** and PE (**f-j**) species distribution in amygdala from HC (**a,f**), people with sPD_>30y_ (**b,g**) and PD-GBA_risk_ (**c,h**) (category (i)), sPD_<10y_ (**d,i**) and PD-GBA_severe_ (**e,j**) (category (ii)). Analysis: 2-way anova with individual comparison mean HC vs each mean of PD group for each species. **(k-r)** Changes in the levels of PS 36:1 (**k,o**), PS 40:6 (**l,p**), PE 36:2 (**m,q**) and PE 40:6 (**n,r**) expressed as mol% of lipid class with GCase activity (**k-n**) or pSer-αS (**o-r**). Analysis: Multiple linear regression of the lipid data, GCase activity and pSer-αS. The black stippled lines represent the 95% confidence interval of the regression model to the data. P-values:t-test of the correlation coefficient of the fitted parameter (GCase activity (**k-n**) or pSer-αS (**o-r**)). \**P* < 0.05, \*\**P* < 0.01, \*\*\**P* < 0.001.

For the SL, the species distribution of SM and Cer were the most affected in sPD and PD-GBA cases compared to HC (**Fig. 4**). 36:1;2 and 42:2;2 were the dominant species with 36:1;2 accounting for ca. 50% of all species in SM and Cer in HC. The relative proportions of SM 36:1;2 and Cer 36:1;2 were decreased by a factor of 2 whereas those of SM 42:2;2 and Cer 42:2;2 were increased by the same factor in sPD_>30y_ and PD-GBA_risk_ cases relative to HC (**Fig. 4 a-c, f-h**). Opposite changes in the percentage levels of these species were observed in sPD_<10y_ and PD-GBA_severe_ cases relative to HC but to a lesser extent (**Fig. 4 a, d-e, f, i-j**). The species distribution of the other SL was affected to a lesser extent (**Supplementary Fig. 8**). For HexCer, the relative proportion of 36:1;2 and 42:2;2 species were affected in the opposite direction to that observed for SM and Cer, with HexCer 36:1;2 being increased in sPD_>30y_ and/or PD-GBA_risk_ cases compared to HC (**Supplementary Fig. 8**). Finally, the relative proportion of 36:1;2 and 42:2;2 diHexCer species were affected differently than those of SM, Cer and HexCer with the percentage levels of 36:1;2 and 42:2;2 species progressively increasing or decreasing, respectively, with increasing disease duration in sPD cases and from PD-GBA_severe_ to PD-GBA_risk_ cases compared to HC (**Supplementary Fig. 8**).

For the GPL, the species distribution of PE and PS were the most altered in sPD and PD-GBA cases compared to HC (**Fig. 5**). The relative proportion of 40:6 species decreased from ca. 30% of all PE or PS in HC to ca. 2-8% in sPD_>30y_ and PD-GBA_risk_ cases but increased to ca. 40-50% in sPD_<10y_ and PD-GBA_severe_ cases. On the contrary the relative proportion of PS 36:1 and PE 36:2 increased in sPD_>30y_ and PD-GBA_risk_ but decreased in sPD_<10y_ and PD-GBA_severe_. The lipid distribution of other GPL were altered to a lesser extent (**Supplementary Fig. 9-12**).

For both SL and GPL, we found that the species for which the relative proportion was increased in sPD_>30y_ and PD-GBA_risk_ (e.g. SM 42:2;2, Cer 42:2;2, PS 36:1 and PE 36:2) correlated positively with pSer-αS and negatively with GCase (**Fig. 4k-r**, **5k-r**). On the contrary, the species for which the relative proportion was decreased in sPD_>30y_ and PD-GBA_risk_ (e.g. SM 36:2;2, Cer 36:2;2, PE and PS 40:6) correlated negatively with pSer-αS and positively with GCase (**Fig. 4k-r**, **5k-r**).

Such changes in species distribution observed in the two categories of people with PD, category (i) (sPD_>30y_ and PD-GBA_risk_) and category (ii) (sPD_<10y_ and PD-GBA_severe_) translated into changes in the average chain length (ChL) and number of double bonds (DB) of the corresponding lipid class. Both the ChL and DB were systematically decreased for most GPL classes, HexCer and diHexCer whereas they were found to be increased for SM and Cer in sPD_>30y_ and PD-GBA_risk_ compared to their values in HC and the opposite was observed for sPD_<10y_ and PD-GBA_severe_ (**Supplementary Fig. 13**).

Altogether, these results show that the relative proportion of specific SL species (36:1;2 and 42:2;2) and GPL species (mainly PS 36:1, PE 36:2 and 40:6 for both PE and PS) are altered in opposite directions in sPD_>30y_ and PD-GBA_risk_ cases (category (i)) on the one hand and sPD_<10y_ and PD-GBA_severe_ cases (category (ii)) on the other hand compared to HC. We observed an increase in the relative proportion of longer, less saturated SM and Cer and shorter, more saturated GPL in sPD_>30y_ and PD-GBA_risk_ cases and the opposite in sPD_<10y_ and PD-GBA_severe_ compared to HC. Finally, we found that the relative proportion of SL and GPL species affected in sPD and PD-GBA cases correlated with both GCase activity and pSer-αS.

## Discussion

Increasing evidence suggests that disruptions in lipid metabolism play a role in the development and spreading of αS pathology in PD ^17–20^. Changes in the levels of specific lipids such as Chol, fatty acids and SL have been observed in association with changes in the levels of total, oligomeric and/or pathological αS in cellular and animal models of PD as well as in tissue samples from people with PD ^17,18^. Mutations in the *GBA* gene, encoding GCase, are, altogether, the most important genetic risk factor for PD and are associated with a decrease in the protein and/or activity levels of GCase, lysosomal dysfunction and/or accumulation of pathological αS and/or the GCase substrate, GlcCer and GlcSph in fibroblasts, CSF and post-mortem brain samples ^18,21,22^. A decrease in GCase activity has been reported not only in samples from people with PD carriers of a *GBA* mutation but also in samples from people with sporadic PD ^18^. Different brain areas are variably affected by decreased GCase activity and/or protein levels, which are associated with increased pathological αS and specific lipid accumulation ^18^.

This study investigates the interplay between GCase, αS, and lipids in amygdala homogenates from HC, people with sPD and varying disease durations and people with PD carrier of a *GBA* mutation. We chose to study amygdala tissues as this brain area is affected by high αS pathology with limited neuronal loss ^54^. Moreover, the amygdala is affected at different stages in the body-first vs brain-first model making this brain area central to the understanding of these two types of PD ^55,56^.

Our combined lipidomic and biochemical analyses of amygdala homogenates from HC, people with sPD and people with PD carriers of a *GBA* mutation show that the levels of GCase activity were decreased whereas the levels of pathological αS, total lipid, specific GPL, Chol, glycerides and SL were increased in sPD_>30y_ and PD-GBA_risk_ (i.e. p.E365K and p.T408M) cases relative to their levels in HC (**Fig. 6**). The levels of lipids altered in sPD_>30y_ and PD-GBA_risk_ amygdala were found to correlate with both GCase activity and pathological αS in two opposite ways: (i) SL, Chol, DAG and specific GPL all correlated positively with pSer-αS and negatively with GCase activity, only below a threshold corresponding to ca. 85% of GCase activity in healthy samples whereas (ii) CL and PE did not correlate with pSer-αS but correlated positively with GCase activity over the whole range of values. The fact that SM, that is abundant in lysosomes^57^, and CL and PE, that are synthetised in mitochondria^57^, correlate in opposite ways with GCase activity, suggests that these two organelles are affected by a decrease in GCase activity likely via different pathways.

**Figure 6:**
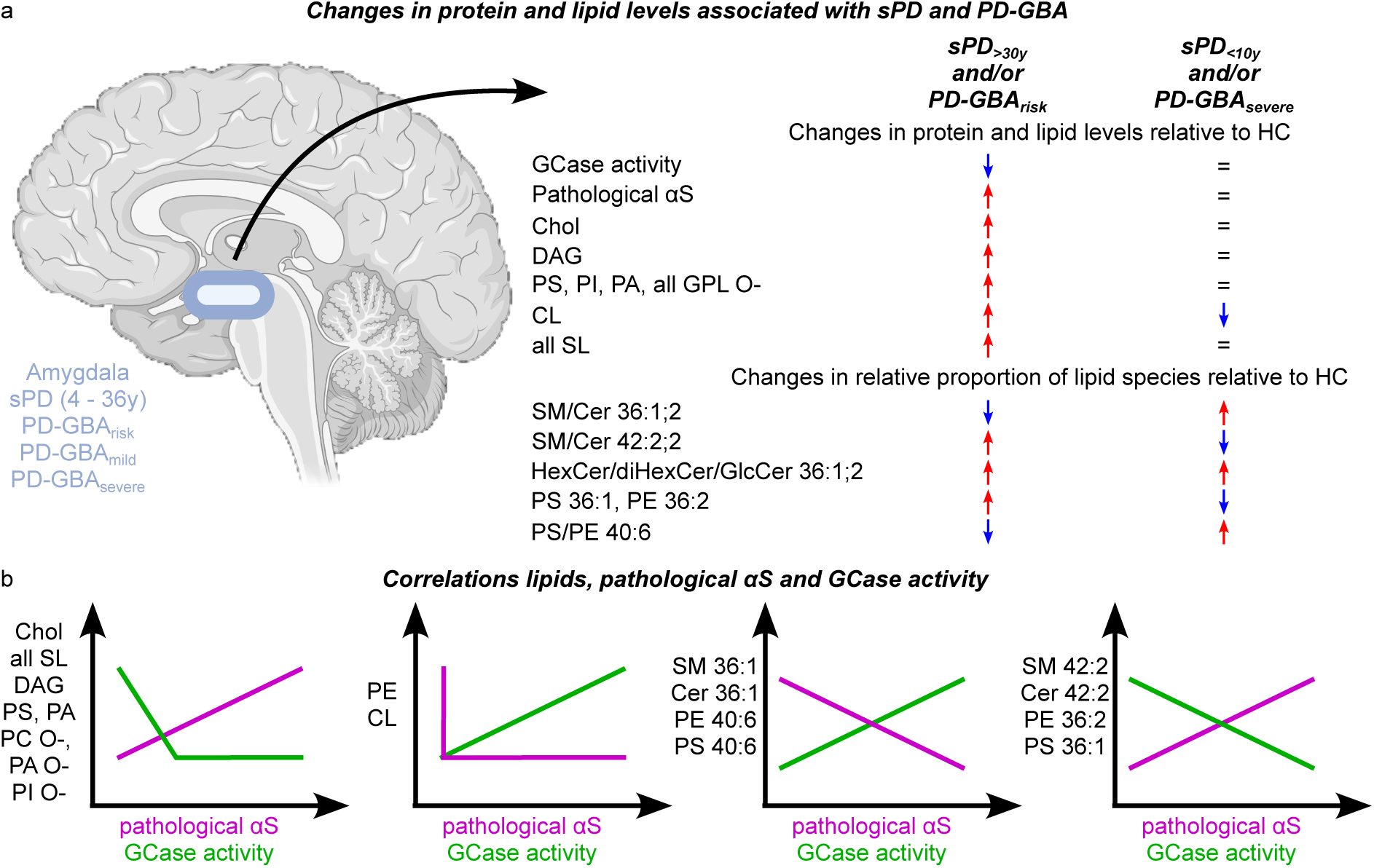
Summary of the main findings of this paper. (**a**) Significant changes in the levels of pathological αS, GCase and main lipids observed in sPD and PD-GBA cases relative to HC. (**b**) Correlation between lipid classes or lipid species and pSer-αS and GCase activity.

The species distribution of SM, Cer, PS and PE were also disrupted in PD amygdala relative to HC. We found that the relative proportion of the same lipid species were altered in these samples but in opposite direction for (i) sPD_>30y_ and PD-GBA_risk_, on the one hand, and (ii) sPD_<10y_ and PD-GBA_severe_ amygdala, on the other hand. In particular, the relative proportions of Cer 36:1;2, SM 36:1;2, PS 40:6 and PE 40:6 were found to decrease whereas the relative proportion of Cer 42:2;2, SM 42:2;2, PS 36:1 and PE 36:2 to increase in sPD_>30y_ and PD-GBA_risk_ amygdala (**Fig. 6**). The opposite was observed sPD_<10y_ and PD-GBA_severe_ amygdala but to a lesser extent (**Fig. 6**). The observed shift from long to short SM and Cer in PD-GBA_severe_ amygdala was also observed in human fibroblasts from people with PD carrier of the p.L483P (severe) *GBA* mutation (PD-GBA_p.L483P_)^32^, suggesting that p.L483P *GBA* mutation is associated with the same lipid changes in fibroblasts and brain samples. A similar shift from long to short chain was observed for SM and Cer in anterior cingulate cortex and for SM in amygdala from people with sPD and disease duration below 30 years ^45,58^.

In addition to Cer, SM, PE and PS, the species distribution of CL was also altered in sPD_>30y_ and PD-GBA_risk_ amygdala relative to HC (**Supplementary Fig. 12**) translating to a decrease in the average ChL and DB of CL (**Supplementary Fig. 13**). The relative proportion of 72:5 or 72:6 CL were increased whereas those of 74:9 CL were decreased in PD_>30y_ and PD-GBA_risk_ amygdala relative to HC (**Supplementary Fig. 12**). Moreover, the relative proportion of 72:5 and 74:9 CL correlated negatively and positively, respectively with GCase activity. CL and PE are commonly associated with mitochondria because they are both synthesised in this organelle, with CL being exclusively synthetised and found in mitochondria and PE being synthetised in mitochondria and in the endoplasmic reticulum. CL and PE are major components of mitochondrial membranes ^57^ and CL is proposed to be involved in mitochondrial morphology, bioenergetics, dynamics, and signalling pathways ^59^. Changes in the levels of brain CL have been proposed to be associated with impaired neuronal function and neurodegeneration ^59^. Our results suggest that disruptions in the levels and properties of CL may be responsible for the reported mitochondrial dysfunction associated with *GBA* mutations in fibroblasts and induced pluripotent stem cell (iPSC)-derived neurons ^60,61^. Moreover, PE and CL are two non-bilayer lipids. Changes in their levels and chain properties can create changes in the properties of the membrane and affect the function of membrane proteins as well as the binding of αS to the membrane and the process of protein-lipid co-assembly into amyloid fibrils. Indeed, the shift from long to short PS was found to lead to faster αS-PS co-aggregation^13^. The observed decrease in the average ChL and DB for both CL and PE observed in sPD_>30y_ and PD-GBA_risk_ amygdala could thus not only be responsible for mitochondrial dysfunction but also for pathological interaction between αS and the mitochondrial membrane.

Correlations between the levels of GCase (activity/protein), αS (pathological/soluble/insoluble) and/or some SL, including GCase substrates GlcCer and GlcSph, SM and Cer, have been identified in fibroblasts and specific brain areas (including ACC, cingulate and frontal cortices, amygdala and Cerebellum) across HC and/or PD samples ^18–20,32,42,45^. Our combined lipidomic and biochemical analyses of HC, sPD and PD-GBA amygdala confirm these findings as we found that not only GlcCer but also all SL and their species composition correlate with both GCase activity and pathological αS. Moreover, our study shows that these correlations were not limited to SL but were also observed for other lipids such as chol, DAG and some GPL. Although unexpected, the correlation between GCase activity and GPLs and their species composition may be explained by an influence of GCase on the cellular fatty acid pool via the lysosomal degradation of SL. The degradation of SL involves GCase followed by the cleavage of the N-acyl chain off the sphingoid backbone by ceramide synthase. Changes in the SL species composition would thus lead to changes in the composition of the fatty acids released upon degradation and as a result to a change in the species composition of GPL that are synthetised via incorporation of these fatty acids. More studies are required to provide the molecular mechanism behind the correlation between GCase, αS and lipids as well as to establish the cause-and-effect relationship among these levels in the context of PD.

The αS Origin site and Connectome (SOC) model proposes that PD can be divided into 2 subtypes, i.e. brain first and body first PD^56^. In people with brain first PD, pathology begins in the central nervous system (often in the amygdala) and then spread to the gut via the vagus nerve. In contrast, in people with body first PD, pathology begins in the peripheral autonomic nervous system and spreads to the brain also via the vagus nerve. The SOC model thus proposes that people with brain first PD would show more pathology in amygdala than people with body first PD. Our results show that the levels of pathological αS, Chol, SL and specific GPL were the highest in amygdala from people with sPD_>30y_ and PD-GBA_risk_ suggesting that sPD_>30y_ and PD-GBA_risk_ could correspond to brain-first PD whereas sPD_<10y_ and PD-GBA_severe_ to body-first PD. This hypothesis suggests that lipid levels and species distribution may be considered or taken into account in the definition of the two subtypes of PD. Characterisation of pathological αS and lipid levels in additional brain areas would be required to confirm this hypothesis.

In conclusion, our combined lipidomic and biochemical analyses of HC, sPD and PD-GBA amygdala show that the levels of GCase activity are decreased whereas those of pathological αS are increased in PD_>30y_ and PD-GBA_risk_ cases. PD and PD-GBA amygdala were also characterised by extensive metabolic remodelling of brain lipids, including increased Chol, DAG, SL and specific GPL and disruptions in the distribution of GPL and SL species (**Fig. 6**). We identified two opposite changes in species distribution for the following groups: (i) a shift towards shorter less unsaturated PE and PS and towards longer more unsaturated SM and Cer for sPD_>30y_ and PD-GBA_risk_ and the opposite for sPD_<10y_ and PD-GBA_severe_. Moreover, the levels of affected lipid classes and relative proportion of species were found to correlate with GCase activity and pathological αS levels when combining all amygdala samples (HC, sPD, PD-GBA) (**Fig. 6**). The fact that we observed opposite lipid changes in PD-GBA_risk_ and PD-GBA_severe_ amygdala suggests that substrate alternating approach aiming at reverting lipid changes associated with a given *GBA* mutation might benefit carriers of *GBA* mutations like *GBA* risk but would result in worse outcome in carriers of *GBA* severe mutations. Our findings thus reinforce the need for patient stratification that may explain previous failures in clinical trials.

## Funding

This work was supported by Lundbeck Foundation (C.G.), Novo Nordisk Foundation (J.E.D., K.S.S., C.G.), Carlsberg Foundation (S.S.M., F.R.M., C.G.), Parkinsonsforeningen (F.R.M., C.G.), Independent Research Fund Denmark (Danmarks Frie Forskningsfond) (K.S., C.G.), Innovation Fund Denmark (Innovationsfonden) under the frame of ERA Permed (F.R.M., C.G.).

## Ethics declarations

### Conflict of interest

The authors have no competing interests to declare that are relevant to the content of this article.

### Human ethics and consent to participate

This study was approved by the London – Central Research Ethics Committee (London REC reference: 23/LO/0044) and the committees on health research ethics in the Capital Region of Denmark (Journal no.: H-21012684). All brain samples were obtained from Queen Square Brain Bank for Neurological Disorders (London, UK) following local research approval and have informed consent from patients and/or their next-of-kind for material usage for medical research. Our research was conducted in accordance with the Declaration of Helsinki.

Clinical trial number: not applicable

## Notes

### Competing Interest Statement

The authors have declared no competing interest.

### Summary of Updates

This version of the manuscript has been revised to include healthy controls and to improve the data analyses.

## References

1. Damier, P., Hirsch, E. C., Agid, Y. & Graybiel, A. M. The substantia nigra of the human brain: II. Patterns of loss of dopamine-containing neurons in Parkinson’s disease. Brain 122, 1437–1448 (1999).

2. Shahmoradian, S. H. et al. Lewy pathology in Parkinson’s disease consists of crowded organelles and lipid membranes. Nat Neurosci 22, 1099–1109 (2019).

3. Spillantini, M. G. et al. α-Synuclein in Lewy bodies. Nature 388, 839–840 (1997).

4. Theillet, F.-X. et al. Structural disorder of monomeric α-synuclein persists in mammalian cells. Nature 530, 45–50 (2016).

5. Burré, J. et al. Properties of native brain α-synuclein. Nature 498, E4–E6 (2013).

6. Fusco, G. et al. Direct observation of the three regions in α-synuclein that determine its membrane-bound behaviour. Nat Commun 5, 3827 (2014).

7. Bendor, J. T., Logan, T. P. & Edwards, R. H. The Function of α-Synuclein. Neuron 79, 1044–1066 (2013).

8. Fink, A. L. The Aggregation and Fibrillation of α-Synuclein. Acc. Chem. Res. 39, 628– 634 (2006).

9. Galvagnion, C. et al. Lipid vesicles trigger α-synuclein aggregation by stimulating primary nucleation. Nat Chem Biol 11, 229–234 (2015).

10. Grey, M. et al. Acceleration of α-Synuclein Aggregation by Exosomes*. Journal of Biological Chemistry 290, 2969–2982 (2015).

11. Martinez, Z., Zhu, M., Han, S. & Fink, A. L. GM1 Specifically Interacts with α-Synuclein and Inhibits Fibrillation. Biochemistry 46, 1868–1877 (2007).

12. Zhu, M. & Fink, A. L. Lipid Binding Inhibits α-Synuclein Fibril Formation*. Journal of Biological Chemistry 278, 16873–16877 (2003).

13. Galvagnion, C. et al. Chemical properties of lipids strongly affect the kinetics of the membrane-induced aggregation of α-synuclein. Proceedings of the National Academy of Sciences 113, 7065–7070 (2016).

14. Galvagnion, C. et al. Lipid Dynamics and Phase Transition within α-Synuclein Amyloid Fibrils. J. Phys. Chem. Lett. 10, 7872–7877 (2019).

15. Hellstrand, E., Nowacka, A., Topgaard, D., Linse, S. & Sparr, E. Membrane Lipid Co-Aggregation with α-Synuclein Fibrils. PLOS ONE 8, e77235 (2013).

16. Galvagnion, C. et al. Structural characterisation of α-synuclein–membrane interactions and the resulting aggregation using small angle scattering. Phys. Chem. Chem. Phys. 26, 10998–11013 (2024).

17. Galvagnion, C. The Role of Lipids Interacting with α-Synuclein in the Pathogenesis of Parkinson’s Disease. Journal of Parkinson’s Disease 7, 433–450 (2017).

18. Muñoz, S. S., Petersen, D., Marlet, F. R., Kücükköse, E. & Galvagnion, C. The interplay between Glucocerebrosidase, α-synuclein and lipids in human models of Parkinson’s disease. Biophysical Chemistry 273, 106534 (2021).

19. Leyns, C. E. G. et al. Glucocerebrosidase activity and lipid levels are related to protein pathologies in Parkinson’s disease. npj Parkinsons Dis. 9, 1–14 (2023).

20. Galper, J. et al. Lipid pathway dysfunction is prevalent in patients with Parkinson’s disease. Brain 145, 3472–3487 (2022).

21. Gegg, M. E. & Schapira, A. H. V. The role of glucocerebrosidase in Parkinson disease pathogenesis. The FEBS Journal 285, 3591–3603 (2018).

22. Schapira, A. H. V., Chiasserini, D., Beccari, T. & Parnetti, L. Glucocerebrosidase in Parkinson’s disease: Insights into pathogenesis and prospects for treatment. Movement Disorders 31, 830–835 (2016).

23. Sidransky, E. et al. Multi-center analysis of glucocerebrosidase mutations in Parkinson disease. N Engl J Med 361, 1651–1661 (2009).

24. Gan-Or, Z. et al. Differential effects of severe vs mild GBA mutations on Parkinson disease. Neurology 84, 880–887 (2015).

25. Parlar, S. C., Grenn, F. P., Kim, J. J., Baluwendraat, C. & Gan-Or, Z. Classification of GBA1 variants in Parkinson’s disease; the GBA1-PD browser. Mov Disord 38, 489–495 (2023).

26. Mallett, V. et al. GBA p.T369M substitution in Parkinson disease: Polymorphism or association? A meta-analysis. Neurol Genet 2, e104 (2016).

27. Duran, R. et al. The Glucocerobrosidase E326K Variant Predisposes to Parkinson’s Disease, But Does Not Cause Gaucher’s Disease. Mov Disord 28, 232–236 (2013).

28. Schöndorf, D. C. et al. iPSC-derived neurons from GBA1-associated Parkinson’s disease patients show autophagic defects and impaired calcium homeostasis. Nat Commun 5, 4028 (2014).

29. Ambrosi, G. et al. Ambroxol-induced rescue of defective glucocerebrosidase is associated with increased LIMP-2 and saposin C levels in *GBA1* mutant Parkinson’s disease cells. Neurobiology of Disease 82, 235–242 (2015).

30. Sanchez-Martinez, A. et al. Parkinson disease-linked GBA mutation effects reversed by molecular chaperones in human cell and fly models. Sci Rep 6, 31380 (2016).

31. Collins, L. M., Drouin-Ouellet, J., Kuan, W.-L., Cox, T. & Barker, R. A. Dermal fibroblasts from patients with Parkinson’s disease have normal GCase activity and autophagy compared to patients with PD and GBA mutations. Preprint at 10.12688/f1000research.12090.2 (2018).

32. Galvagnion, C. et al. Sphingolipid changes in Parkinson L444P GBA mutation fibroblasts promote α-synuclein aggregation. Brain 145, 1038–1051 (2022).

33. Sun, Y. et al. Properties of Neurons Derived from Induced Pluripotent Stem Cells of Gaucher Disease Type 2 Patient Fibroblasts: Potential Role in Neuropathology. PLOS ONE 10, e0118771 (2015).

34. Yang, S., Gegg, M., Chau, D. & Schapira, A. Glucocerebrosidase activity, cathepsin D and monomeric α-synuclein interactions in a stem cell derived neuronal model of a PD associated *GBA1* mutation. Neurobiology of Disease 134, 104620 (2020).

35. Yang, S.-Y., Beavan, M., Chau, K.-Y., Taanman, J.-W. & Schapira, A. H. V. A Human Neural Crest Stem Cell-Derived Dopaminergic Neuronal Model Recapitulates Biochemical Abnormalities in *GBA1* Mutation Carriers. Stem Cell Reports 8, 728–742 (2017).

36. Fernandes, H. J. R. et al. ER Stress and Autophagic Perturbations Lead to Elevated Extracellular α-Synuclein in *GBA-N370S* Parkinson’s iPSC-Derived Dopamine Neurons. Stem Cell Reports 6, 342–356 (2016).

37. Woodard, C. M. et al. iPSC-Derived Dopamine Neurons Reveal Differences between Monozygotic Twins Discordant for Parkinson’s Disease. Cell Reports 9, 1173–1182 (2014).

38. Burbulla, L. F. et al. A modulator of wild-type glucocerebrosidase improves pathogenic phenotypes in dopaminergic neuronal models of Parkinson’s disease. Science Translational Medicine 11, eaau6870 (2019).

39. Rocha, E. M. et al. Progressive decline of glucocerebrosidase in aging and Parkinson’s disease. Annals of Clinical and Translational Neurology 2, 433–438 (2015).

40. Huebecker, M. et al. Reduced sphingolipid hydrolase activities, substrate accumulation and ganglioside decline in Parkinson’s disease. Molecular Neurodegeneration 14, 40 (2019).

41. Gegg, M. E. et al. Glucocerebrosidase deficiency in substantia nigra of parkinson disease brains. Annals of Neurology 72, 455–463 (2012).

42. Murphy, K. E. et al. Reduced glucocerebrosidase is associated with increased α-synuclein in sporadic Parkinson’s disease. Brain 137, 834–848 (2014).

43. Gündner, A. L. et al. Path mediation analysis reveals GBA impacts Lewy body disease status by increasing α-synuclein levels. Neurobiology of Disease 121, 205–213 (2019).

44. Blumenreich, S. et al. Elevation of gangliosides in four brain regions from Parkinson’s disease patients with a GBA mutation. npj Parkinsons Dis. 8, 1–11 (2022).

45. Abbott, S. K. et al. Altered ceramide acyl chain length and ceramide synthase gene expression in Parkinson’s disease. Movement Disorders 29, 518–526 (2014).

46. Nielsen, I. Ø. et al. Comprehensive Evaluation of a Quantitative Shotgun Lipidomics Platform for Mammalian Sample Analysis on a High-Resolution Mass Spectrometer. J. Am. Soc. Mass Spectrom. 31, 894–907 (2020).

47. Pedregosa, F. et al. Scikit-learn: Machine Learning in Python. Journal of Machine Learning Research 12, 2825–2830 (2011).

48. Jekel, C. F. & Venter, G. pwlf: A Python Library for Fitting 1D Continuous Piecewise Linear Functions.

49. Seabold, S. & Perktold, J. Statsmodels: Econometric and Statistical Modeling with Python. scipy (2010) doi:10.25080/Majora-92bf1922-011.

50. Braak, H. et al. Amygdala pathology in Parkinson’s disease. Acta Neuropathol 88, 493– 500 (1994).

51. Harding, A. J., Stimson, E., Henderson, J. M. & Halliday, G. M. Clinical correlates of selective pathology in the amygdala of patients with Parkinson’s disease. Brain 125, 2431–2445 (2002).

52. Sorrentino, Z. A. et al. Unique α-synuclein pathology within the amygdala in Lewy body dementia: implications for disease initiation and progression. Acta Neuropathologica Communications 7, 142 (2019).

53. Dasgupta, S. & Ray, S. K. Diverse Biological Functions of Sphingolipids in the CNS: Ceramide and Sphingosine Regulate Myelination in Developing Brain but Stimulate Demyelination during Pathogenesis of Multiple Sclerosis. J Neurol Psychol 5, 10.13188/2332-3469.1000035 (2017).

54. Fu, Y. et al. Increased unsaturated lipids underlie lipid peroxidation in synucleinopathy brain. acta neuropathol commun 10, 165 (2022).

55. Borghammer, P. The brain-first vs. body-first model of Parkinson’s disease with comparison to alternative models. J Neural Transm 130, 737–753 (2023).

56. Horsager, J. & Borghammer, P. Brain-first vs. body-first Parkinson’s disease: An update on recent evidence. Parkinsonism Relat Disord 122, 106101 (2024).

57. van Meer, G., Voelker, D. R. & Feigenson, G. W. Membrane lipids: where they are and how they behave. Nat Rev Mol Cell Biol 9, 112–124 (2008).

58. Fu, Y. et al. A protective role of ABCA5 in response to elevated sphingomyelin levels in Parkinson’s disease. npj Parkinsons Dis. 10, 1–12 (2024).

59. Falabella, M., Vernon, H. J., Hanna, M. G., Claypool, S. M. & Pitceathly, R. D. S. Cardiolipin, Mitochondria, and Neurological Disease. Trends in Endocrinology & Metabolism 32, 224–237 (2021).

60. Schöndorf, D. C. et al. The NAD+ Precursor Nicotinamide Riboside Rescues Mitochondrial Defects and Neuronal Loss in iPSC and Fly Models of Parkinson’s Disease. Cell Reports 23, 2976–2988 (2018).

61. García-Sanz, P. et al. N370S-GBA1 mutation causes lysosomal cholesterol accumulation in Parkinson’s disease. Movement Disorders 32, 1409–1422 (2017).

