## Supplementary Information for "Shared and Distinct Lipid Profiles in amygdala from Sporadic and GBA-associated Parkinson’s Diseases"

### Supplementary Tables:

**Table S1: Internal lipid standards**

| Lipid class | Formula | Source | ID | Amount added (pmol) |
|---|---|---|---|---|
| CE | CE-D7 15:0 | Avanti Polar Lipids | 700144 | 29.3 |
| Ceramide (Cer) | Cer 18:1;2/12:0;0 | Avanti Polar Lipids | 860512 | 31.3 |
| Ceramide-1-phosphate (CerP) | CerP 18:1;2/12:0;0 | Avanti Polar Lipids | 860531 | 25.0 |
| Cholesterol | Chol-D4 | QMX | D-6359 | 356.9 |
| CL | CL 14:0/14:0/14:0/14:0 | Avanti Polar Lipids | 710332 | 72.2 |
| DAG | DAG 12:0/12:0 | Avanti Polar Lipids | 800812 | 45.6 |
| diHexCer | diHexCer 18:1;2/17:0;0 | Avanti Polar Lipids | 860595 | 23.2 |
| FA | FA 16:0-D4 | TRC-Canada | P145502 | 37.5 |
| GlcCer | GlcCer 18:1;2/12:0;0 | Avanti Polar Lipids | 860543 | 28.9 |
| GalCer | GalCer 18:1;2/12:0;0 | Avanti Polar Lipids | 860544 | 28.9 |
| GM1-3 | GM3 18:1;2/18:0;0-D3 | Larodan | 71-1107 | 37.5 |
| HexCer | GlcCer 18:1;2/12:0;0 and GalCer 18:1;2/12:0;0 | Avanti Polar Lipids | 860544 and 860544 | 57.8 |
| HexSph | GluSph-$^{13}C_6$ 18:1;2 | Larodan | 78-4018 | 25.0 |
| LPA and LPA O- | LPA 17:0 | Avanti Polar Lipids | 11067 | 31.1 |
| LPC and LPC O- | LPC 12:0 | Avanti Polar Lipids | 855475 | 25.0 |
| LPE and LPE O- | LPE 13:0 | Avanti Polar Lipids | 110696 | 30.8 |
| LPG and LPG O- | LPG 17:1 | Avanti Polar Lipids | 858127 | 30.2 |
| LPI and LPI O- | LPI 13:0 | Avanti Polar Lipids | 110716 | 22.9 |
| LPS and LPS O- | LPS 17:1 | Avanti Polar Lipids | 858141 | 14.3 |
| PA and PA O- | PA 12:0/12:0 | Avanti Polar Lipids | 840635 | 28.3 |
| PC and PC O- | PC 12:0/12:0 | Avanti Polar Lipids | 850335 | 35.9 |
| PE and PE O- | PE 12:0/12:0 | Avanti Polar Lipids | 850702 | 39.5 |
| PG and PG O- | PG 12:0/12:0 | Avanti Polar Lipids | 840435 | 29.3 |
| PI and PI O- | PI 8:0/8:0 | Avanti Polar Lipids | 850181 | 28.4 |
| PS and PS O- | PS 12:0/12:0 | Avanti Polar Lipids | 840038 | 25.4 |
| SHexCer | SHexCer 18:1;2/12:0;0 | Avanti Polar Lipids | 860573 | 35.8 |
| SM | SM 18:1;2/12:0;0 | Avanti Polar Lipids | 860583 | 21.3 |
| TAG | TAG 17:0/17:0/17:0 | Larodan | 33-1700 | 907.5 |
| triHexCer | triHexCer18:1;2/17:0;0 | Larodan | 56-1061 | 51.2 |

**Table S2: Ions and neutral losses used for lipid identifications**

| Lipid class | Mode | MS1: Precursor ion | MS2: Fragment ion | MS2: Neutral loss | MS2: m/z | MS2: Species specifics |
|---|---|---|---|---|---|---|
| PC, PC O– | POS | [M+H]+ | [Phosphorylcholine + H]+ | | 184,0733 | All |
| SM | POS | [M+H]+ | [Phosphorylcholine + H]+ | | 184,0733 | All |
| CE | POS | [M+NH4]+ | [Chol – NH3 – H2O] + | | 369,3516 | All |
| Chol | POS (SIM/tPRM) | [M+NH4]+ | [Chol – NH3 – H2O] + | | 369,3516 | All |
| Cer, diHexCer, triHexCer | POS | [M+H] + | [LCB + H – H2O] + | * | * | All |
| | | | [LCB + H – 2H2O] + | * | * | All |
| DAG | POS | [M+NH4]+ | | [Fatty acid – H + NH4] | Delta, * | All |
| CerP | NEG | [M–H]– | [Phosphoric acid – H – H2O] – | | 78,959 | All |
| PS, PS O–, LPS, LPS O– | NEG | [M–H] – | [Glycerophosphate – H – H2O]– | | 152,9958 | All |
| | | | | [C3H5NO2] | Delta, 87.032 | All |
| | | | [Fatty acid – H]– | | * | PS, PS O–, LPS |
| | | | [Fatty acid O– – H]– | | * | PS O– |
| LPC, LPC O– | NEG | [M–H]– | [Fatty acid – H]– | | * | LPC |
| PE, PE O–, LPE, LPE O– | NEG | [M–H]– | [Ethanolaminephosphate – H – H2O]– | | 196,038 | All |
| | | | [Fatty acid – H]– | | * | PE, PE O–, LPE |
| | | | [Fatty acid O– – H]– | | * | PE O– |
| PI, PI O–, LPI, LPI O– | NEG | [M–H]– | [Glycerophosphate – H – H2O]– | | 152,9958 | All |
| | | | [Inositolphosphate – H – H2O]– | | 241,0119 | All |
| | | | [Fatty acid – H]– | | * | PI, PI O–, LPI |
| | | | [Fatty acid O– – H]– | | * | PI O– |

| | | | | | | |
|---|---|---|---|---|---|---|
| PA, PA O–, LPA, LPA O– | NEG | [M–H]– | [Glycerophosphate – H – H2O]– | | 152,9958 | All |
| | | | [Fatty acid – H]– | | * | PA, PA O–, LPA |
| | | | [Fatty acid O– – H]– | | * | PA O– |
| BMP/PG, LPG, LPG O– | NEG | [M–H]– | [Glycerophosphate – H – H2O]– | | 152,9958 | All |
| | | | [Fatty acid – H]– | | * | BMP/PG, LPG |
| SHexCer | NEG | $[M–H]^-$ | $[HO_4S]^-$ | | 96,9601 | All |
| GM3, GM2, GM1 | NEG | $[M–H]^-$ | $[NeuAc – H]^-$ | | 290,0864 | All |
| HexCer | NEG | $[M–H]^-$ | | | | |
| CL | NEG | $[M–2H]^{2-}$ | | | | |

**Table S3: p- values for the correlation between lipid classes levels and Age at death (AAD), post-mortem intervals (PMI) and GCase protein**

| Lipid classes | p-value Correlation AAD | p-value Correlation PMI | p-value Correlation Gcase protein |
|---|---|---|---|
| Chol | 0.17839525 | 0.52432206 | 0.6599 |
| CE | 0.9435812 | 0.50663735 | 0.5825 |
| PC | 0.47061815 | 0.51809067 | 0.5036 |
| PE | 0.62458127 | 0.38576319 | 0.6823 |
| PS | 0.25541216 | 0.56448426 | 0.9016 |
| PI | 0.25086594 | 0.59811461 | **0.041** |
| CL | 0.93388287 | 0.34536807 | 0.0561 |
| PA | 0.08396106 | 0.5505275 | 0.5744 |
| PG | 0.12411875 | 0.66618106 | 0.8075 |
| PE O- | 0.51919918 | 0.58926976 | 0.8738 |
| PC O- | 0.26580919 | 0.59299232 | 0.9588 |
| PA O- | 0.12113249 | 0.67074027 | 0.9871 |
| PI O- | 0.17157385 | 0.48027854 | 0.8373 |
| LPE | 0.45495416 | 0.62983294 | 0.8096 |
| LPC | 0.88876262 | 0.61951129 | 0.8375 |
| LPI | 0.35242755 | 0.63184374 | 0.0699 |
| LPS | 0.39383326 | 0.40755099 | 0.3153 |
| LPA | 0.71585575 | 0.54283404 | 0.73 |
| LPG | 0.2694298 | 0.56374412 | 0.4074 |
| LPE O- | 0.23219399 | 0.47664434 | 0.4341 |
| LPC O- | 0.29706024 | 0.5695234 | 0.4884 |
| SM | 0.17817164 | 0.61858316 | 0.8156 |
| HexCer | 0.33449789 | 0.9391401 | 0.6957 |
| Cer | 0.29826009 | 0.44566353 | 0.9295 |
| SHexCer | 0.38484163 | 0.68509497 | 0.5235 |
| diHexCer | 0.22973199 | 0.40628756 | 0.78 |
| GM3 | 0.58624786 | 0.80578126 | 0.871 |
| CerP | 0.27225894 | 0.9156796 | 0.7792 |
| DAG | 0.24101939 | 0.95743081 | 0.3754 |
| TAG | 0.18698245 | 0.51016484 | **0.0002** |

### Supplementary Figures

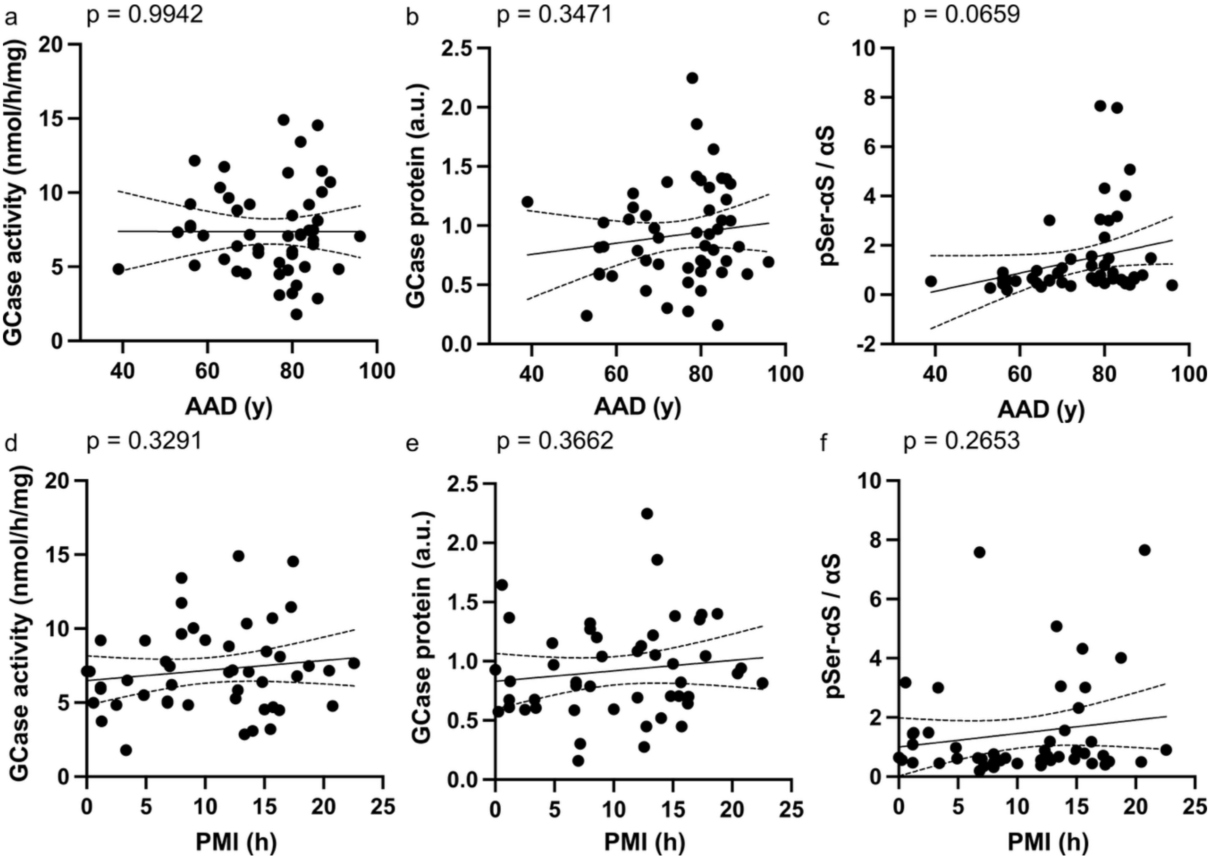


**Figure S1: GCase activity, GCase protein and pSer-αS levels do not correlate with age at death (AAD) and post-mortem intervals (PMI).** (**a-f**) Variation in the levels of GCase activity (**a,d**), GCase protein (**b,e**) and pSer-αS (**c,f**) with age at death (AAD) (**a-c**) or post-mortem intervals (PMI) (**d-f**) among all cases (HC, sPD and PD-GBA). n = 50 (AAD), n = 48 (PMI).

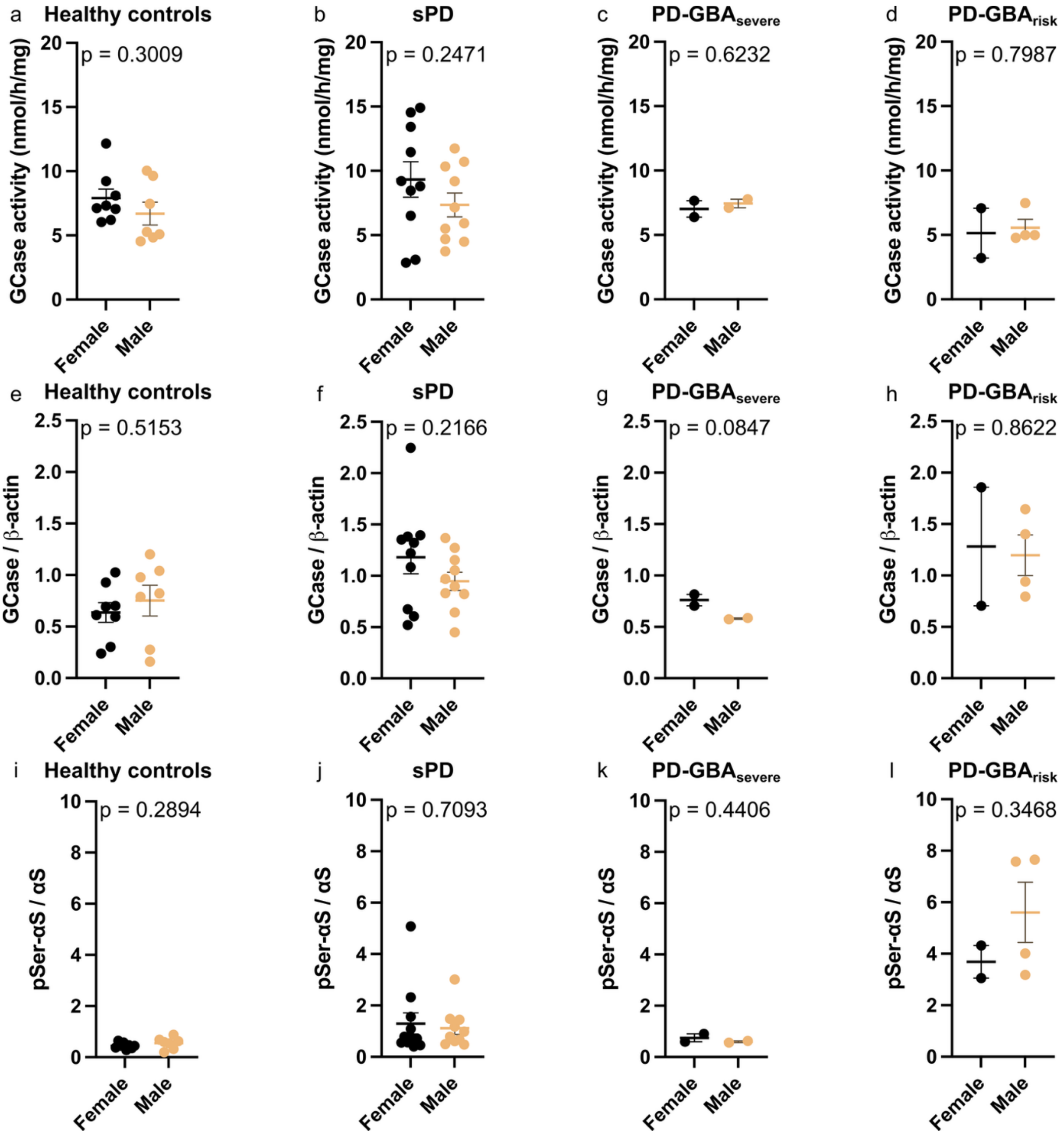


**Figure S2: GCase activity, GCase protein and pSer-αS levels do not differ between female and male cases.** (**a-l**) Levels of GCase activity (**a-d**), GCase protein (**e-h**) and pSer-αS (**i-l**) in female and male cases for the HC (n = 15, F:M = 8:7), sPD (n = 20, F:M = 10:10), PD-GBA$_{severe}$ (n = 4, F:M = 2:2) and PD-GBA$_{risk}$ (n = 5, F:M = 2:4) groups. t-test comparison between the mean of female and that of the male cases in each of the above mentioned 4 groups.

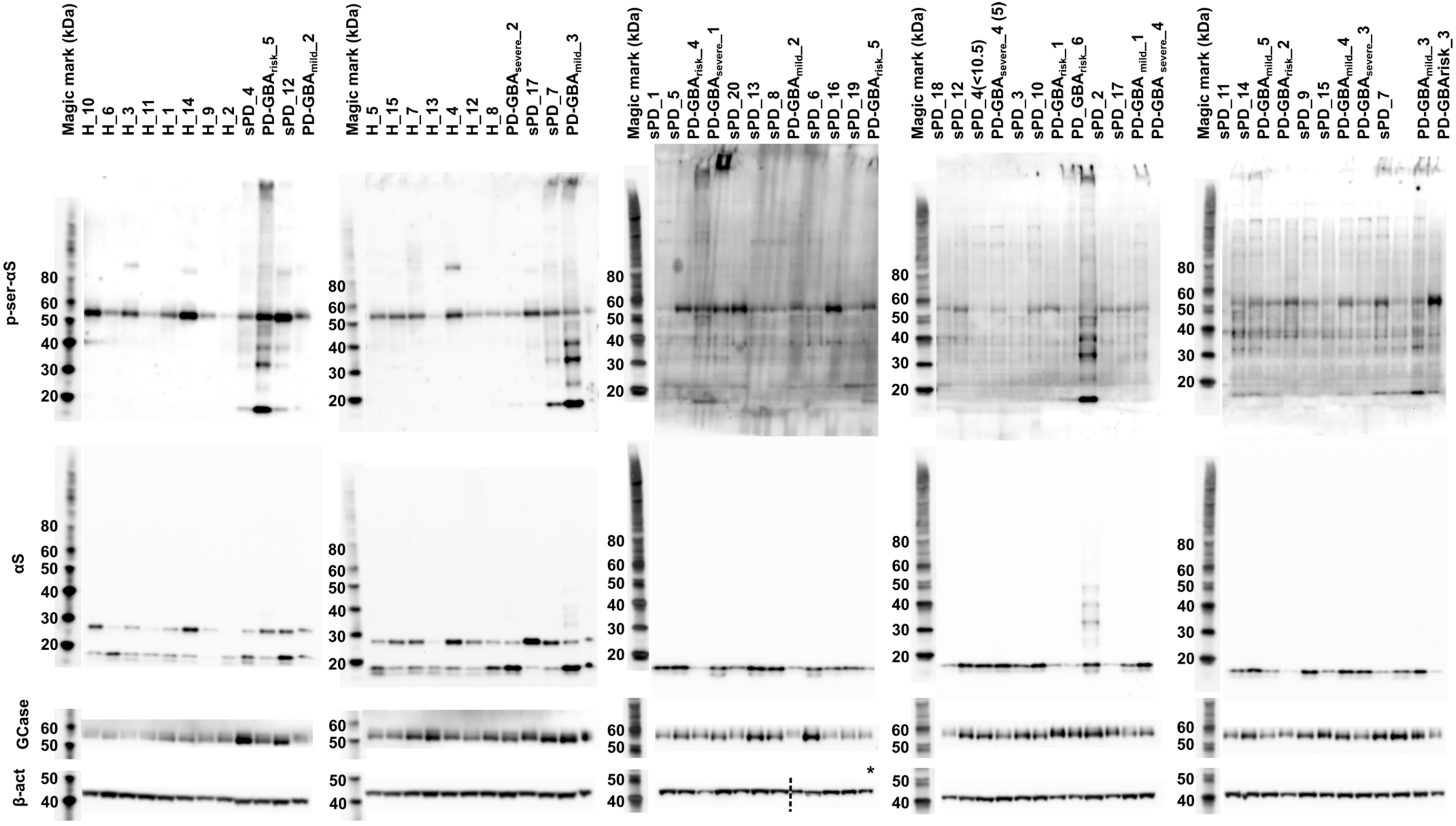
Magic mark (kDa)
H_10
H_6
H_3
H_11
H_1
H_14
H_9
H_2
sPD_4
PD-GBA$_{risk}$_5
sPD_12
PD-GBA$_{mild}$_2
Magic mark (kDa)
H_5
H_15
H_7
H_13
H_4
H_12
H_8
PD-GBA$_{severe}$_2
sPD_17
sPD_7
PD-GBA$_{mild}$_3
Magic mark (kDa)
sPD_1
sPD_5
PD-GBA$_{risk}$_4
PD-GBA$_{severe}$_1
sPD_20
sPD_13
sPD_8
PD-GBA$_{mild}$_2
sPD_6
sPD_16
sPD_19
PD-GBA$_{risk}$_5
Magic mark (kDa)
sPD_18
sPD_12
sPD_4(<10.5)
PD-GBA$_{severe}$_4 (5)
sPD_3
sPD_10
PD-GBA$_{risk}$_1
PD_GBA$_{risk}$_6
sPD_2
sPD_17
PD-GBA $_{mild}$_1
PD-GBA $_{severe}$_4
Magic mark (kDa)
sPD_11
sPD_14
PD-GBA$_{mild}$_5
PD-GBA$_{risk}$_2
sPD_9
sPD_15
PD-GBA$_{mild}$_4
PD-GBA$_{severe}$_3
sPD_7
PD-GBA$_{mild}$_3
PD-GBArisk_3
p-ser-αS
αS
GCase
β-act
80
60
50
40
30
20
*

**Figure S3: Western Blots showing immunoreactivity for human GCase, β-actin, total αS and pathological αS (one replicate per homogenate) in HC, sPD and PD-GBA amygdala homogenates.** For each target protein two images of the same membrane acquired with different exposure times are shown side by side; one image showing the ladder (left) where intensity is adjusted to that of the ladder, the same image showing the samples where the intensity is adjusted to that of the samples bands (right). * This membrane was accidently cut in such a way that the β-actin bands for the last five samples were on the part of the membrane used for GCase staining. Therefore, the two parts of this membrane were together stripped and reprobed for β-actin. The two parts of this membrane are shown side by side in this figure for illustrative purpose.

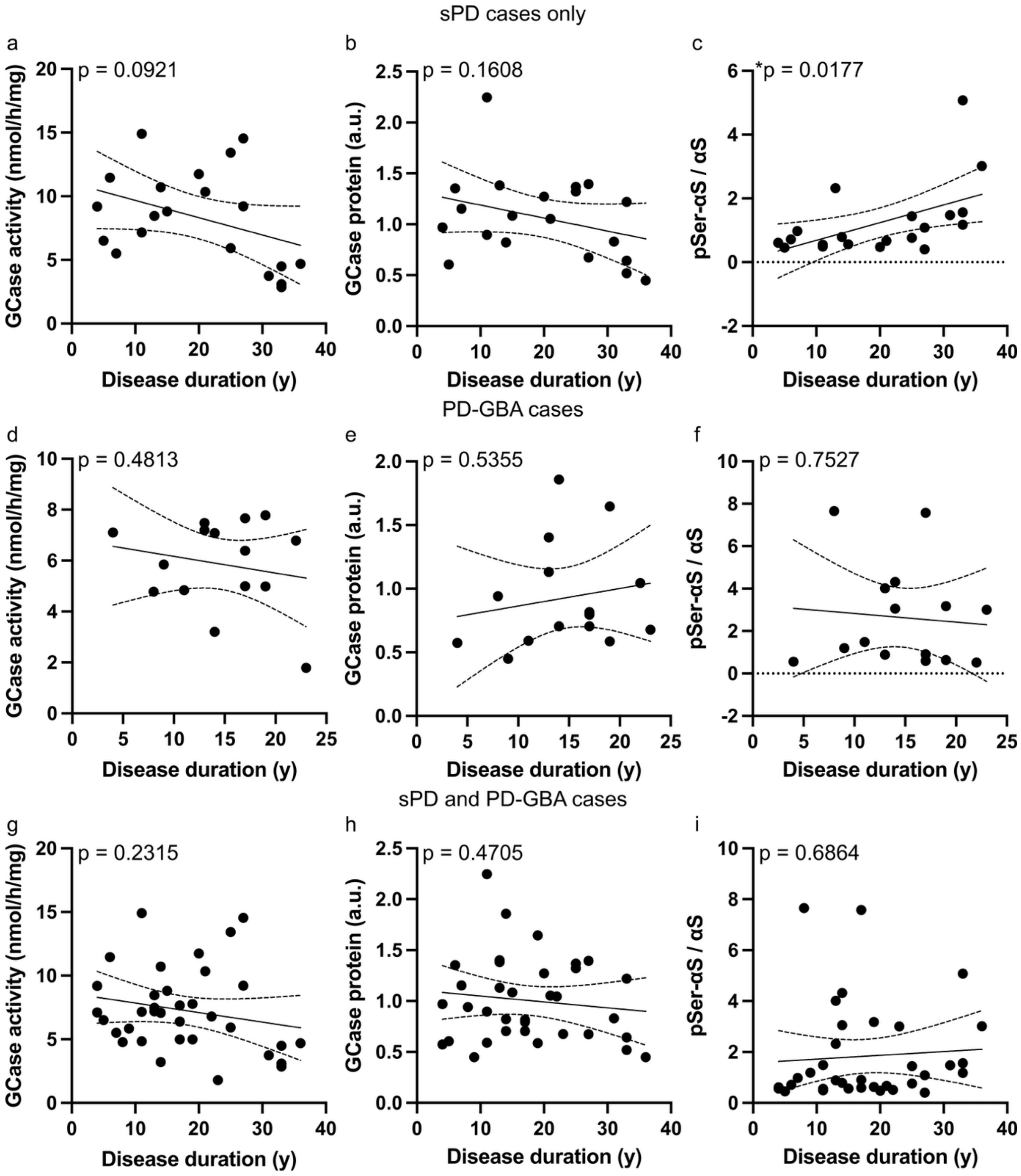


**Figure S4: GCase activity and protein levels do not correlate with disease duration in PD and PD-GBA amygdala but pathological αS only correlate with disease duration in sPD amygdala.** (**a-i**) Variation in GCase activity (**a,d,g**), GCase protein (**b,e,h**) and pSer-αS (**c,f,i**) with disease duration for sPD amygdala only (n = 20, **a-c**), PD-GBA amygdala only (n = 15, **d-f**) or all PD amygdala (n = 35, **g-i**). *$P$ <0.05

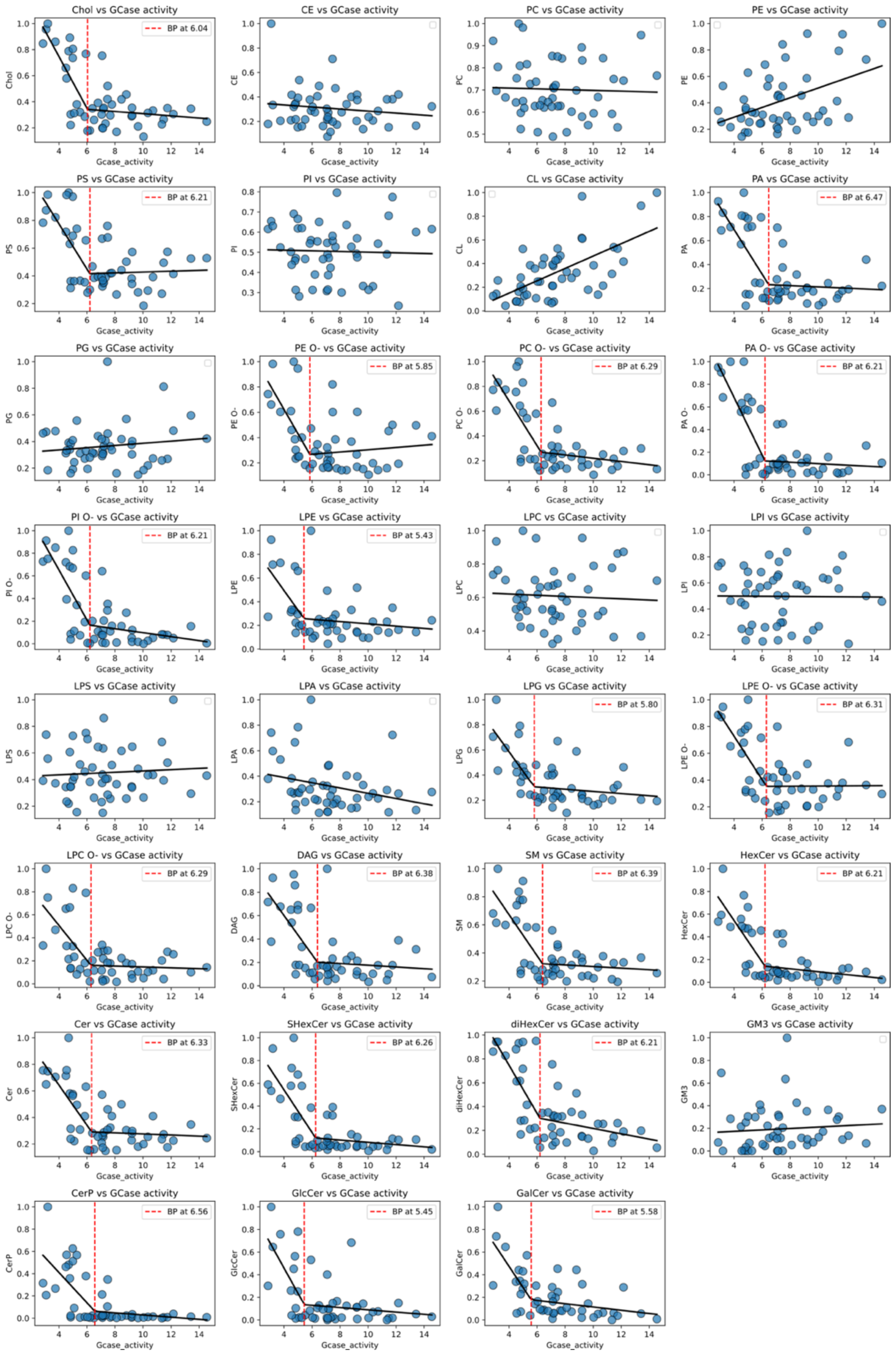


**Figure S5: Normalised levels of lipid classes fitted to a piecewise linear model to estimate the value of GCase activity below which levels correlate with GCase activity.** For each lipid class, lipid levels were normalized to the maximum value, and fitted using the piecewise linear model described in the materials and methods. The black lines show the fitted line, and the red stippled lines show the calculated breakpoint. In plots with no stippled lines and only one linear fit, no breakpoint was detected by the model.

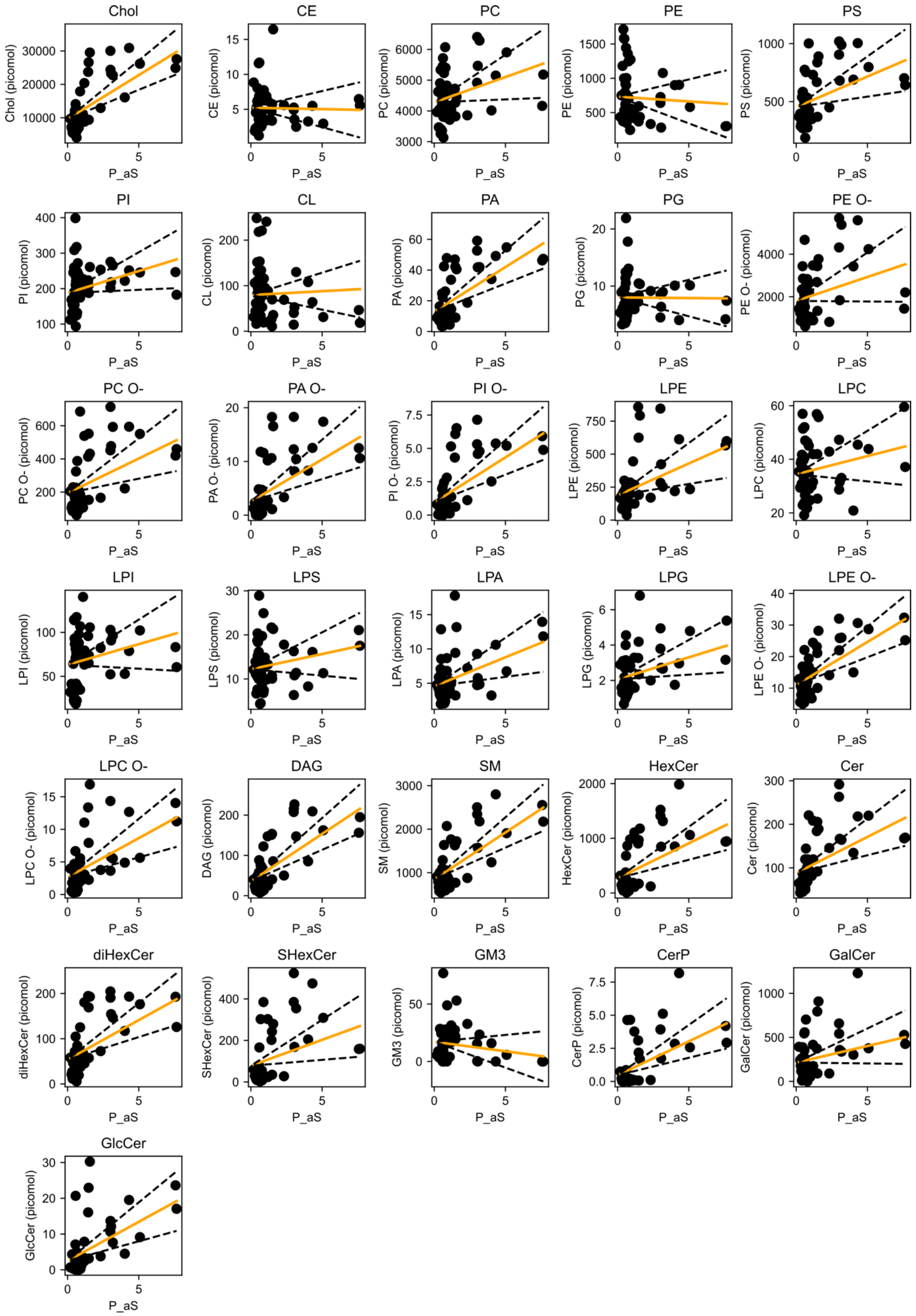


**Figure S6: Changes in the levels of lipid classes expressed in pmol with pSer-αS.** Lipid levels are plotted against pSer-αS values (black dots). The data are fitted to a multiple linear regression model (orange line) adjusted for the contribution of GCase activity, with the confidence interval of the fit shown as black stippled lines.

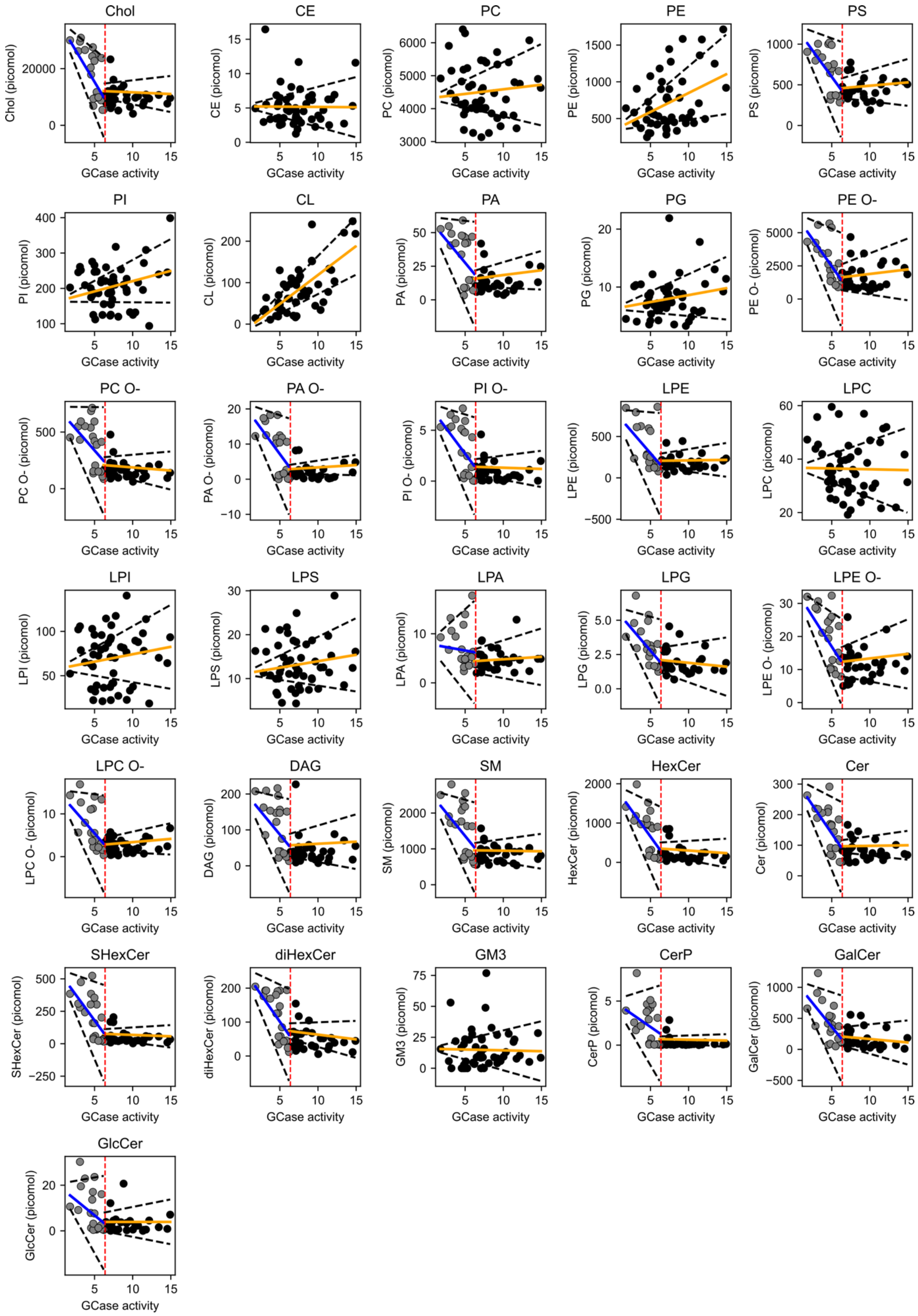


**Figure S7: Changes in the levels of lipid classes expressed in pmol with GCase activity.** Lipid levels are plotted against GCase activity (black dots). The data are fitted to a multiple linear regression model (orange line) adjusted for the contribution of pSer-αS, with the confidence interval of the fit shown as black stippled lines.

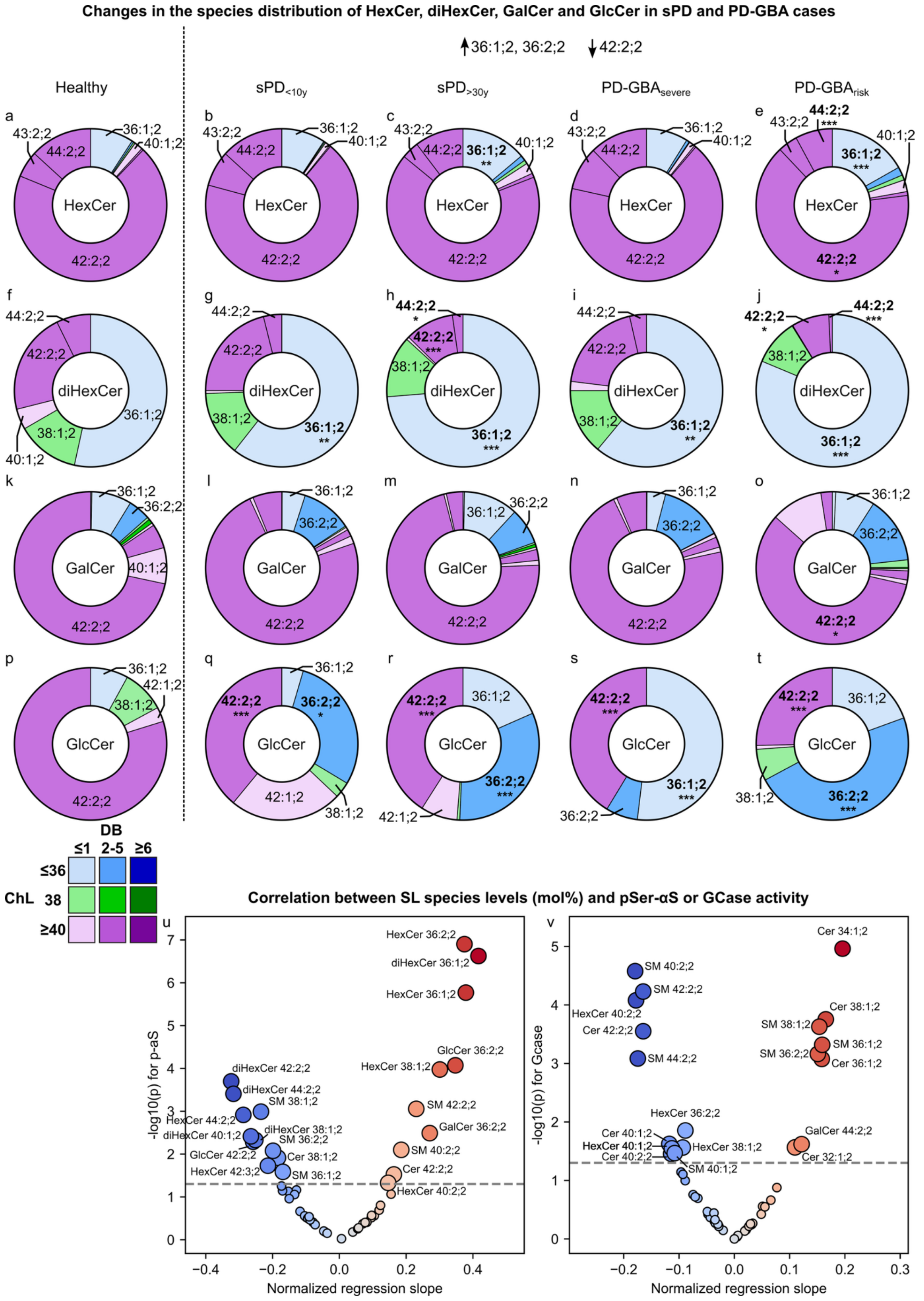


**Figure S8: The levels of specific HexCer, diHexCer, GalCer and GlcCer species are affected in sPD and PD-GBA amygdala homogenates.** **(a-t)** Species distribution of HexCer (**a-e**), diHexCer (**f-l**), GalCer (**k-o**) and GlcCer (**p-t**) in HC (**a,f,k,p**), $sPD_{<10y}$ (**b,g,l,q**), $sPD_{>30y}$ (**c,h,m,r**), PD-$GBA_{severe}$ (**d,i,n,s**), PD-$GBA_{risk}$ (**e,j,o,t**) amygdala. Analysis: 2-way anova with individual comparison mean HC vs each mean of PD group for each species. **(u-v)** Volcano-plot for the multiple linear regression between SL species and pSer-αS adjusted for the contribution of GCase activity (**u**) and GCase activity adjusted for the contribution of pSer-αS (**v**). The scatter dots are coloured by regression slope and annotated if *P*-value is below a 0.05 threshold indicated by the stippled line. *$P < 0.05$, **$P < 0.01$, ***$P < 0.001$.

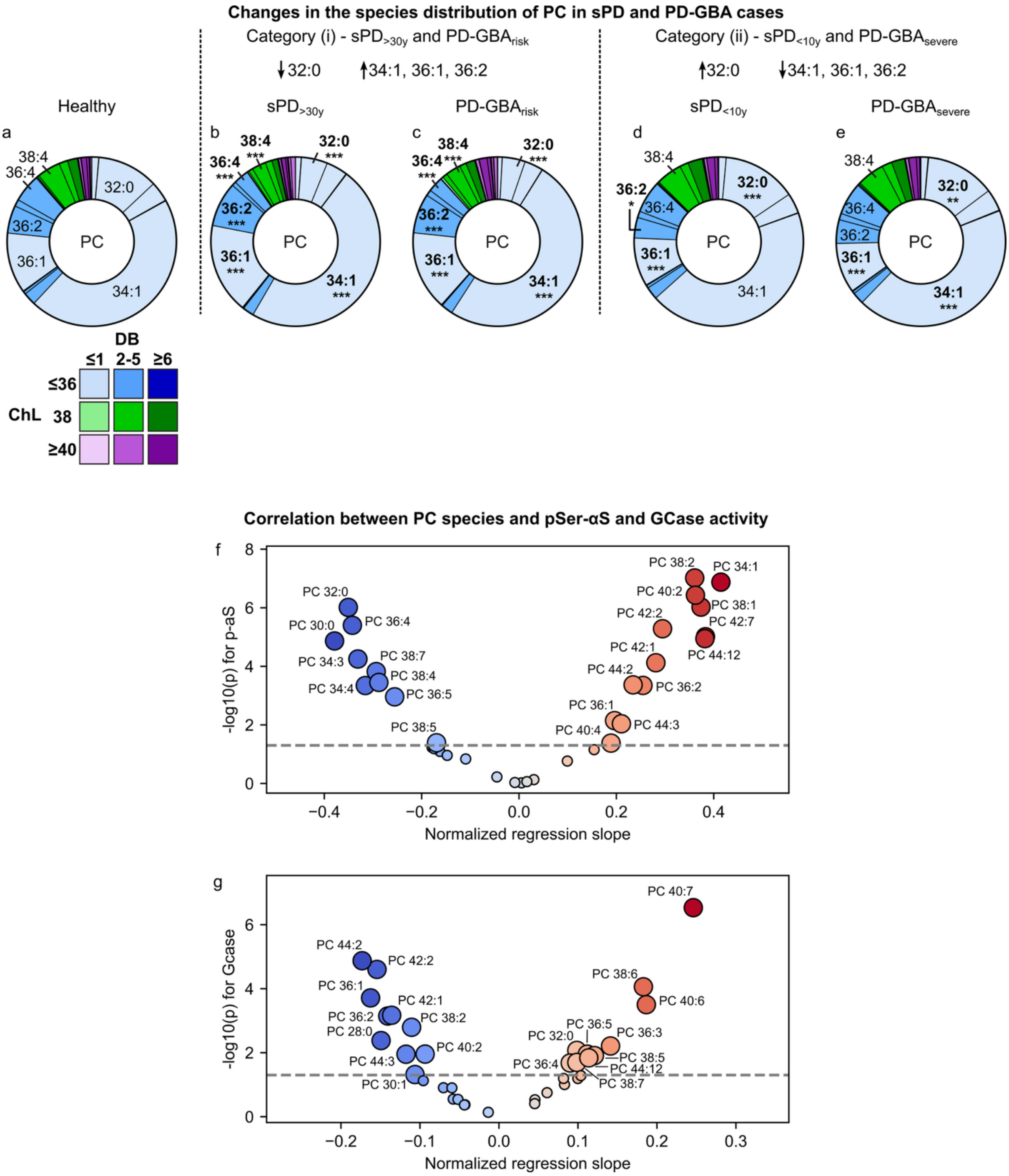


**Figure S9: Changes in the levels of PC species with PD and correlation between their levels and those of pSer-αS and GCase activity.** (**a-e**) PC species distribution in HC (**a**), sPD$_{>30y}$ (**b**) and PD-GBA$_{risk}$ (**c**) (category (i)), sPD$_{<10y}$ (**d**) and PD-GBA$_{severe}$ (**e**) (category (ii)) amygdala. Analysis: 2-way anova with individual comparison mean HC vs each mean of PD group for each species. **(f-g)** Volcano-plot for the multiple linear regression between PC species and pSer-αS adjusted for the contribution of GCase activity (**f**) and GCase activity adjusted for the contribution of pSer-αS (**g**). The scatter dots are coloured by regression slope and annotated if *P*-value is below a 0.05 threshold indicated by the stippled line. *$P < 0.05$, **$P < 0.01$, ***$P < 0.001$.

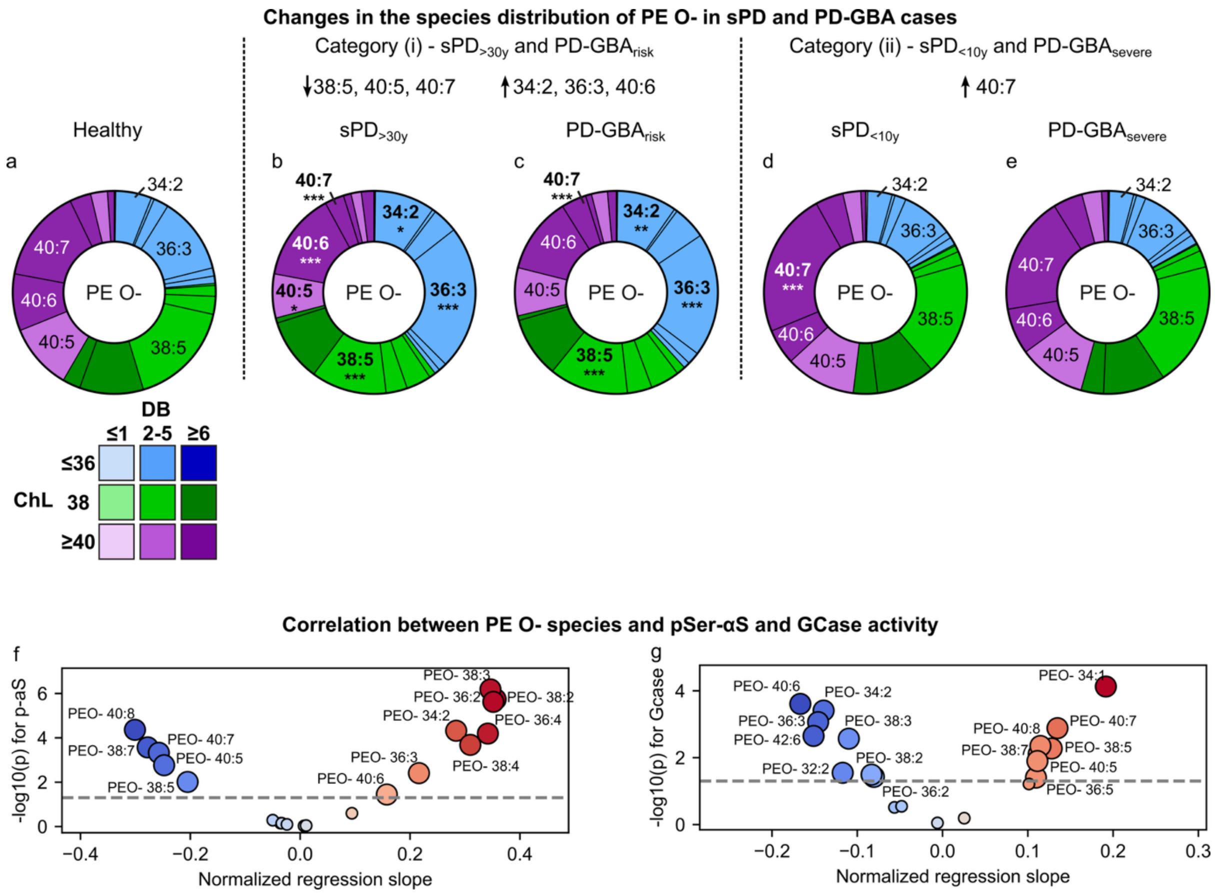


**Figure S10: Changes in the levels of PE O- species with PD and correlation between their levels and those of pSer-αS and GCase activity.** (**a-e**) PE O- species distribution in HC (**a**), $sPD_{>30y}$ (**b**) and $PD\text{-}GBA_{risk}$ (**c**) (category (i)), $sPD_{<10y}$ (**d**) and $PD\text{-}GBA_{severe}$ (**e**) (category (ii)) amygdala. Analysis: 2-way anova with individual comparison mean HC vs each mean of PD group for each species. **(f-g)** Volcano-plot for the multiple linear regression between PE O-species and pSer-αS adjusted for the contribution of GCase activity (**f**) and GCase activity adjusted for the contribution of pSer-αS (**g**). The scatter dots are coloured by regression slope and annotated if *P*-value is below a 0.05 threshold indicated by the stippled line. *$P < 0.05$, **$P < 0.01$, ***$P < 0.001$.

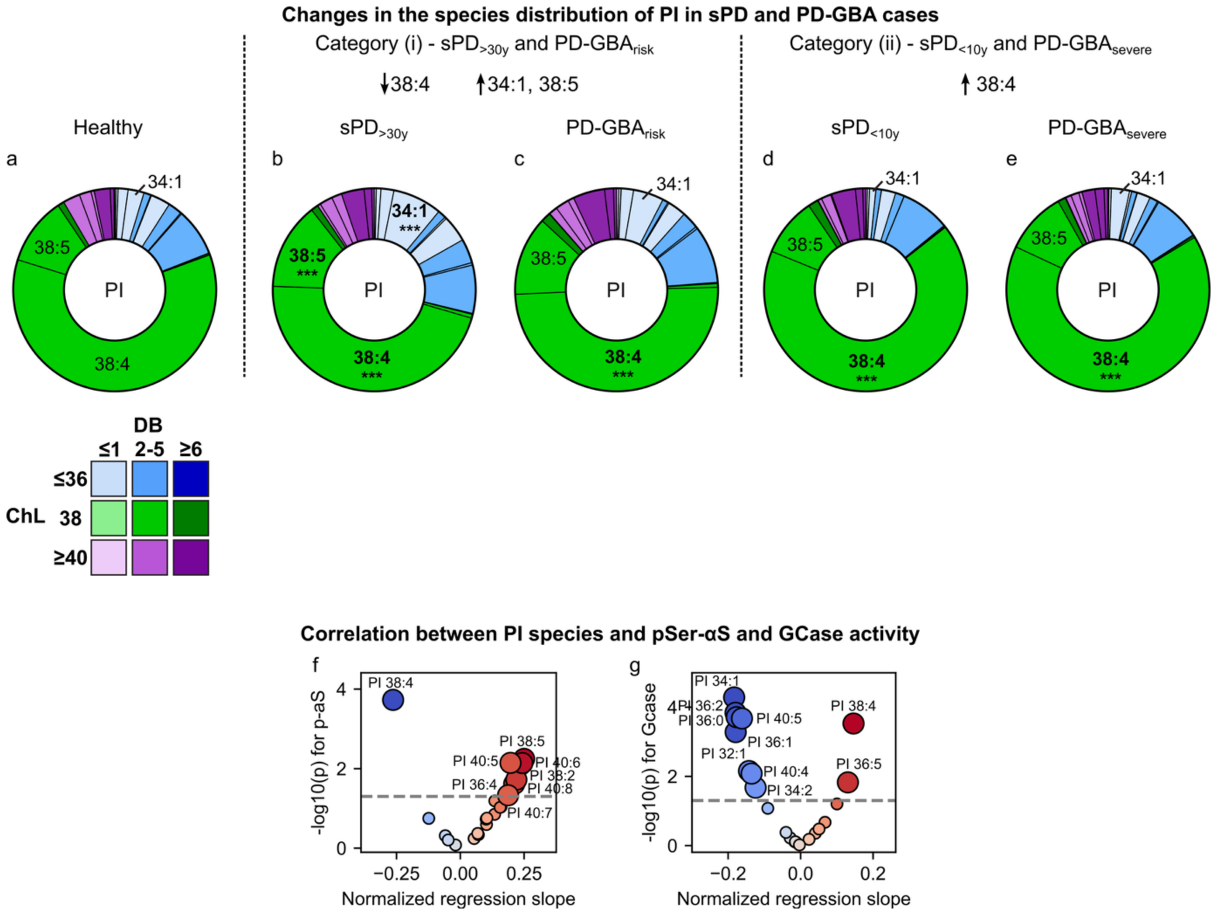


**Figure S11: Changes in the levels of PI species with PD and correlation between their levels and those of pSer-αS and GCase activity.** (**a-e**) PI species distribution in HC (**a**), sPD$_{>30y}$ (**b**) and PD-GBA$_{risk}$ (**c**) (category (i)), sPD$_{<10y}$ (**d**) and PD-GBA$_{severe}$ (**e**) (category (ii)) amygdala. Analysis: 2-way anova with individual comparison mean HC vs each mean of PD group for each species. **(f-g)** Volcano-plot for the multiple linear regression between PI species and pSer-αS adjusted for the contribution of GCase activity (**f**) and GCase activity adjusted for the contribution of pSer-αS (**g**). The scatter dots are coloured by regression slope and annotated if *P*-value is below a 0.05 threshold indicated by the stippled line. *$P < 0.05$, **$P < 0.01$, ***$P < 0.001$.

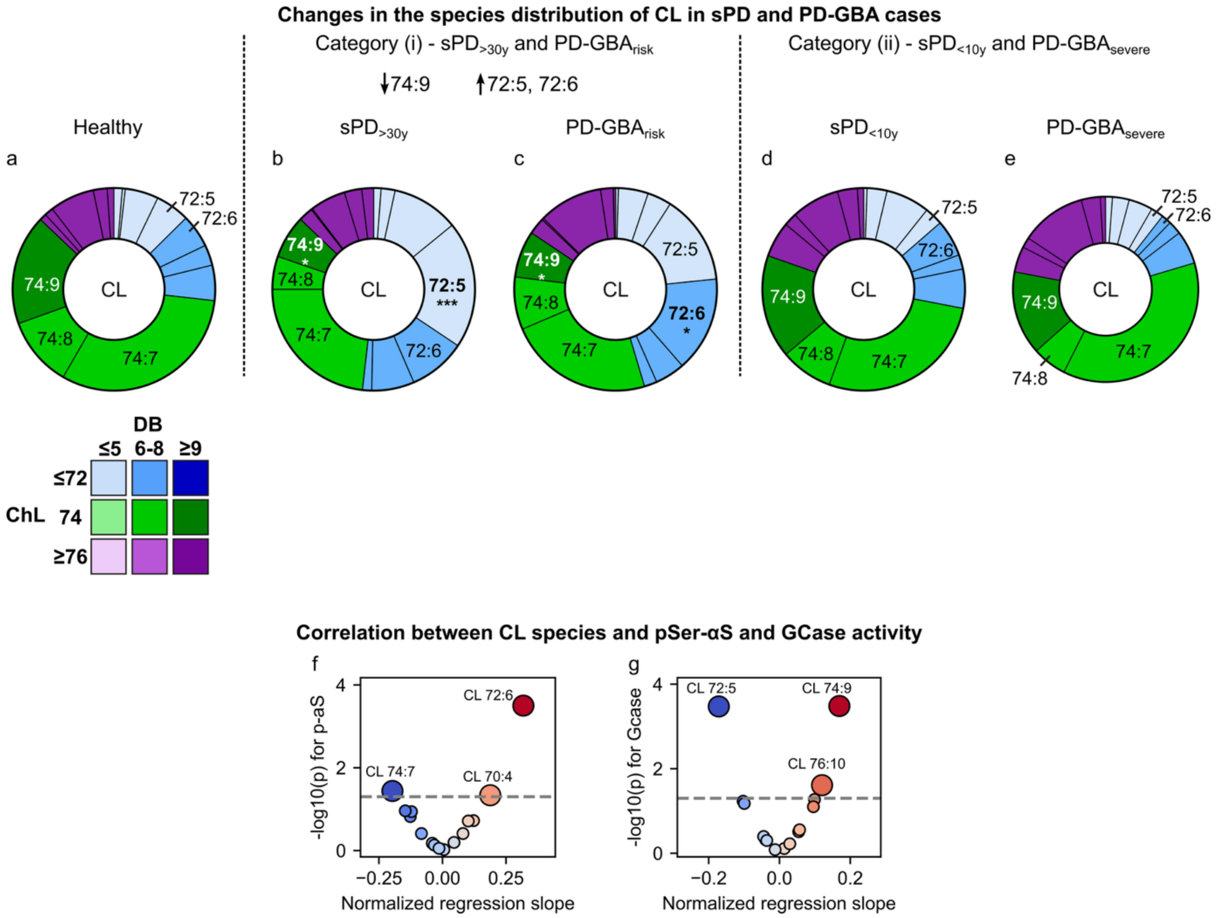


**Figure S12: Changes in the levels of CL species with PD and correlation between their levels and those of pSer-αS and GCase activity.** (**a-e**) CL species distribution in HC (**a**), sPD$_{>30y}$ (**b**) and PD-GBA$_{risk}$ (**c**) (category (i)), sPD$_{<10y}$ (**d**) and PD-GBA$_{severe}$ (**e**) (category (ii)), amygdala. Analysis: 2-way anova with individual comparison mean HC vs each mean of PD group for each species. **(f-g)** Volcano-plot for the multiple linear regression between CL species and pSer-αS adjusted for the contribution of GCase activity (**f**) and GCase activity adjusted for the contribution of pSer-αS (**g**). The scatter dots are coloured by regression slope and annotated if *P*-value is below a 0.05 threshold indicated by the stippled line. **P* < 0.05, ***P* < 0.01, ****P* < 0.001.

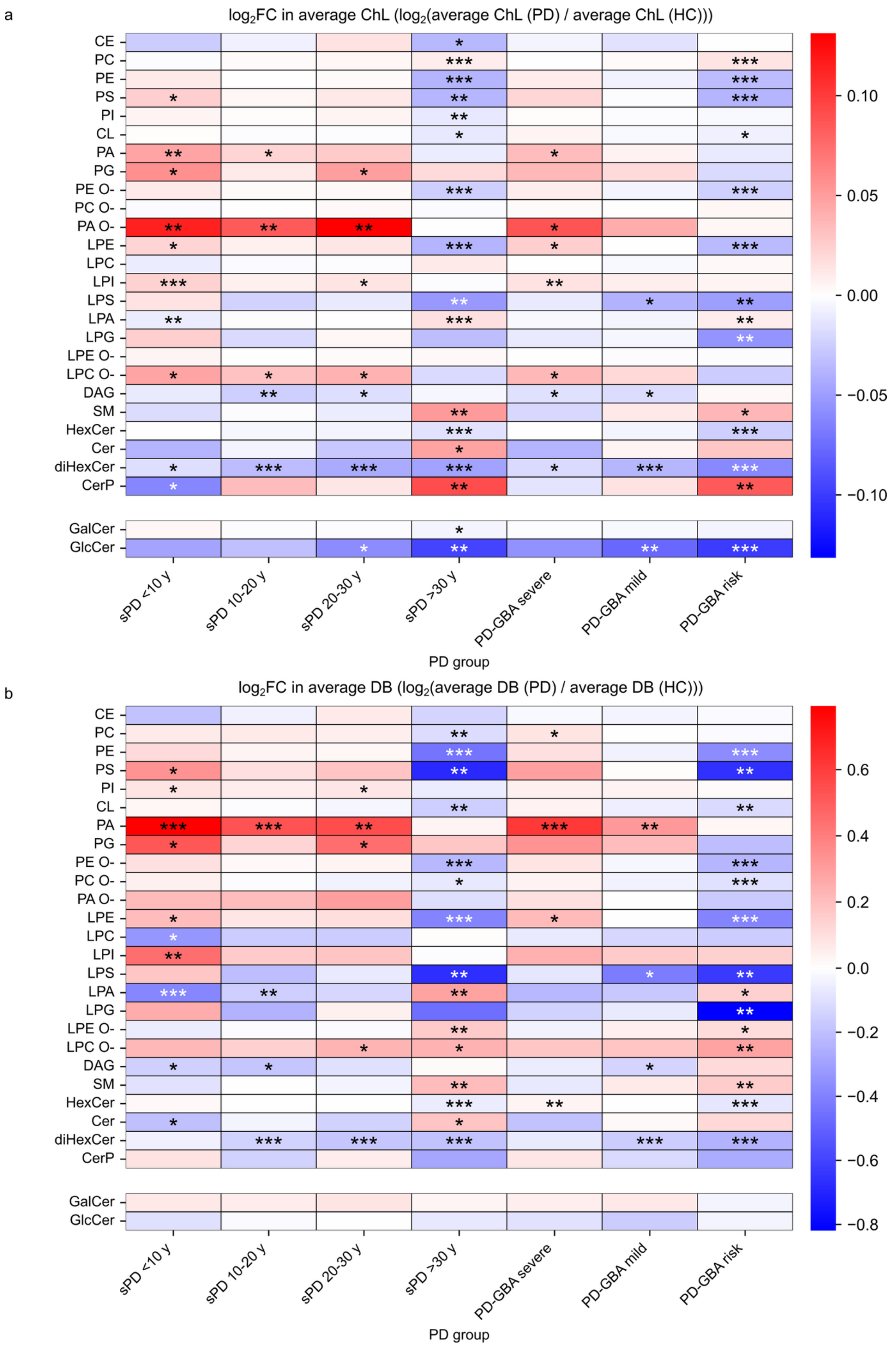
a
log₂FC in average ChL (log₂(average ChL (PD) / average ChL (HC)))
CE
PC
PE
PS
PI
CL
PA
PG
PE O-
PC O-
PA O-
LPE
LPC
LPI
LPS
LPA
LPG
LPE O-
LPC O-
DAG
SM
HexCer
Cer
diHexCer
CerP
GalCer
GlcCer
sPD <10 y
sPD 10-20 y
sPD 20-30 y
sPD >30 y
PD-GBA severe
PD-GBA mild
PD-GBA risk
PD group
0.10
0.05
0.00
−0.05
−0.10
b
log₂FC in average DB (log₂(average DB (PD) / average DB (HC)))
CE
PC
PE
PS
PI
CL
PA
PG
PE O-
PC O-
PA O-
LPE
LPC
LPI
LPS
LPA
LPG
LPE O-
LPC O-
DAG
SM
HexCer
Cer
diHexCer
CerP
GalCer
GlcCer
sPD <10 y
sPD 10-20 y
sPD 20-30 y
sPD >30 y
PD-GBA severe
PD-GBA mild
PD-GBA risk
PD group
0.6
0.4
0.2
0.0
−0.2
−0.4
−0.6
−0.8

**Figure S13: Changes in the average ChL and DB of lipid classes in sPD and PD-GBA cases relative to HC. (a-b)** Heatmaps showing the log2FC in the average chain-length (**a**) and average amount of double bonds (**b**) in each lipid class in sPD and PD-GBA cases relative to that in HC. The stars indicate the p-value for the t-test comparison between the mean of the class level in sPD or PD-GBA cases and that in HC. *$P < 0.05$, **$P < 0.01$, ***$P < 0.001$.
